# Robustness of Selection and Timing Inference under Model Variation in Population Genetics

**DOI:** 10.1101/2025.01.08.631974

**Authors:** Javier Escabi, Sahand Hormoz

## Abstract

In population genetics, accurately inferring the selection coefficient and the time of onset of advantageous mutations from genetic data is fundamental for understanding evolutionary processes. Here, we investigate how mismatches between the true evolutionary process and the inference model—specifically in the reproductive variance (*σ*^2^) and the number of generations (*L*)—affect the posterior distributions of the selection coefficient and the time of onset. Using the Kolmogorov forward and backward equations, we model the stochastic dynamics of gene frequencies under selection and drift. We show that while the posterior distribution of the selection coefficient remains unaffected by changes in *σ*^2^ and *L*, this invariance does not apply to the time of onset. By framing the problem as a first passage time issue, we derive explicit expressions for the offsets in the posterior mean and variance of the time of onset that result from incorrect assumptions about *σ*^2^ and *L*. Our analysis reveals that these offsets are related to the mean and variance of the first passage time required for the allele frequency to reach a certain threshold, starting from an initial frequency determined by the model parameters. Under the assumption of a uniform prior for the time of onset, we find that the offset in the inferred mean is given by the difference in the effective generation duration (Δ = 1*/σ*^2^) between the true process and the inference model. We validate our theoretical findings through simulations, demonstrating that the empirical offsets closely match our predictions. Furthermore, we generalize our results to accommodate non-uniform prior distributions, such as exponential priors, and provide numerical methods for calculating offsets under arbitrary priors. Stochastic fluctuations due to genetic drift, which are influenced by the reproductive variance and generational structure, can introduce significant biases in the posterior distribution of time of onset of advantageous mutations. By quantifying these biases, our framework enables more accurate adjustments to inferences drawn from genetic data, thereby enhancing our understanding of evolutionary dynamics and improving the reliability of population genetic analyses.

## 2 Introduction

Population genetics models of drift and selection have been used in modeling real-world populations with different biological details, for example, variation in offspring number, timing between generations, population structure, and sexual reproduction. These models are used for inference because they are known to be robust. One way to define robustness is to consider the behavior of these models under certain limits. For example, many distinct models including the Wright-Fisher model [1, 2], the Moran model [3], and the Cannings models [4, 5] share the same diffusion limit: population size *N* → ∞, selection coefficient *s* → 0, such that *Ns* → *C*, where *C* is a nonzero constant. In this limit, the probability distribution of gene frequency at time *t >* 0 conditioned on an initial frequency converges to a universal description by the Kolmogorov forward equation under an effective population size [6, 7, 8, 9]. This ensures a robustness where the time-course of gene frequencies for the same initial conditions are identical across many different models as long as the effective population size is the same [9]. A similar type of robustness has been defined for the ancestral relationships of genes in addition to their relative frequencies in the population. In the same diffusion limit and in the neutral case, the gene genealogy of a finite random sample of genes backwards in time converges to the Kingman coalescent under an appropriate scaling of time according to the effective population size [10, 11, 12]. General conditions for convergence to the Kingman coalescent have been extended to include a changing population size, non-exchangeable offspring number distributions, selfing and sexual reproduction, etc., through a generalized time-scale transformation [10, 11, 12, 13, 14, 15, 16, 17]. From the perspective of inference, these results guarantee a robustness of the Kingman coalescent in a different sense: only an effective population size, defined as the ratio of true population size *N* to a model specific reproductive variance *σ*^2^, can be inferred from an observed set of coalescence times. However, inference of population size *N* itself is not robust because it requires knowledge of *σ*^2^.

The idea of robustness historically has been centered around inferences of effective population size to uncover demographic histories that include population bottlenecks, population structure, and population growth [18, 19, 20, 21, 22, 23, 24], reviewed in [25, 26]. Inference of selection has become of interest more recently due to the drop in cost and advances in modern sequencing technologies, for example, inferences of population growth rates in human populations due to the increased statistical power to observe rare genetic variants [27, 28, 29, 30, 31], reviewed in [32], inferences of selection and timing of driver mutations in blood cancers from single-cell whole genome sequencing experiments [33, 34], and inferences of selection in the drosophila genome [35, 36, 37]. On the theoretical side, models of selection have been extensively studied, including balancing selection [38], the ancestral selection graph [39], multiple merger models [40, 41, 42, 43], and selective sweeps [43, 44, 45, 46, 47, 48, 49, 50, 51], reviewed in [52, 53]. However, robustness in the context of 1) inference of selection and 2) inference of the time at which the selectively advantageous mutation arose has received surprisingly little attention. This analysis is particularly important when selection is weak, for example, in the case of inferring selection and timing of the driver mutations of blood cancers which undergo random fluctuations in population size in their early history [33, 34]. Importantly, the fluctuations in the trajectories of the population size of the selectively advantageous mutation created by random drift introduces uncertainty when inferring these two quantities from an observed set of coalescence times. This makes the analysis of the robustness of the inference of these two quantities subtly different from robustness of the coalescent statistics under an effective population size in the classical sense.

We will now pose the central question of this paper: suppose we have a coalescent tree from a population under selection and infer a posterior distribution for a selection coefficient and the time at which the selectively advantageous mutation arose using a model. How will the inferred distributions change if the model were to change?

We can consider other forms of data as well. For example, we can consider the site-frequency spectrum of a sample or time-course measurements of gene frequencies. In all these cases, the results/equations of the paper will apply. Throughout the paper, we will consider coalescent trees since it adds an additional layer of complexity, but will point out how the results are a more general property of population genetics models and apply to other forms of data as well.

We can frame the question more precisely. To do this, we will first define a class of models that include selection. We will begin by defining a modified version of the Wright-Fisher model with selection [1, 2], and then generalize the process into a class of Cannings models with selection [4, 5, 54]. The goal is to model a single mutation with a selective advantage conditional on fixation that begins with a population size of one, although these results can be generalized to multiple competing or subclonal mutations. We will then frame the inference part of the question more precisely after the model definitions.

## 3 Model definitions

We will begin by defining the Wright-Fisher model with selection. The model begins with a total population of *N* identical individuals (or genes), where *N* remains fixed over time. At each generation, the individuals at the current generation give birth to a new generation of *N* individuals and immediately die off. Each individual in the new generation can descend from any individual in the previous generation with equal probability. This process is iterated until at some generation one individual acquires a mutation that gives it a selective advantage *s*. We will refer to *s* as the selection coefficient. From this point on, the probabilities of descent change. Each individual in the new generation descends from wild-type individuals in the previous generation with probability *p*, and from mutant individuals in the previous generation with probability (1 + *s*)*p*. If *n* is the number of mutant individuals in the previous generation, then *p* can be derived from the condition that probabilities must sum to 1:

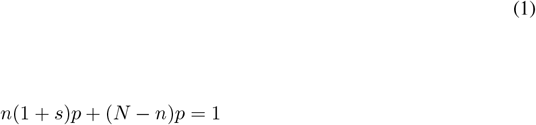

We then define *L* to be the total number of generations of the Wright-Fisher process, and *g* to be the number of generations from the time at which the mutant individual arose all the way to the *L*^*th*^ generation at which the process terminates. Lastly, we define *a* to be the age of the population, or the amount of time (for example in years) that has elapsed from the first generation to the *L*^*th*^. Note that the number of generations per unit time is then 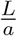, assuming that the duration of each generation remains the same over time.

We are interested in generalizing the Wright-Fisher model with selection in the Cannings sense [1, 2, 4, 5]. A rigorous treatment and definition of the Cannings models with selection and examples of these models can be found in [54, 55]. We will take a similar approach to [54], and construct the generalized Cannings version of our model for completeness.

We will begin by defining a class of models that agrees with the Wright-Fisher model in terms of the average time-course of gene frequencies under selection, but differs in terms of the fluctuations in offspring number. To do this, there are a set of general characteristics of the Wright-Fisher model that we will retain while allowing other characteristics of the model to change. In particular, denote the set of random variables that represents the number of offspring produced by each individual in the Wright-Fisher model at a single generation *t* as 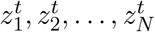. For any fixed *t* before the selectively advantageous mutation arose (prior to generation *L* − *g*), the joint distribution of 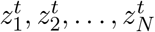 follows a multinomial distribution with parameter values *N*_*i*_ = *N* and 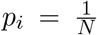, where *i* refers to the *i*^*th*^ individual. Importantly, the offspring numbers at a given generation are exchangeable random variables, where exchangeable means that the marginal distributions are invariant under arbitrary permutations. In addition, 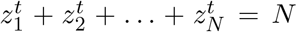 since we assumed the population size is fixed. However, the 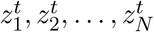 are not exchangeable random variables once the selectively advantageous mutation arises. At this point, the random variables are partitioned into two distinct classes with two different distributions: one class of individuals with a selectively advantageous mutation that on average produces more offspring, and another class of individuals that do not carry the selectively advantageous mutation. Here, the marginal distributions are invariant only if we permute the random variables that belong to the same class, or in other words, exchangeability holds only within each class. To construct the Cannings model, we will retain the condition of exchangeability within each classeq and assume 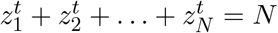, but will allow the distributions of 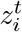 to deviate from the multinomial form at the level of the higher moments beyond the mean. For example, one way to do this is to assume the 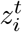 satisfy a multinomial form as in the Wright-Fisher model but draw the *p*_*i*_ from a given distribution [55].

Next we will assume that the variance in the offspring number distribution is a *σ*^2^ value across all the individuals in each generation, and that the mean value of any 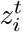 conditional on belonging to a particular class and conditional on the total fraction of individuals belonging to that class (to account for the decreasing effect of the selection coefficient when approaching saturation), matches the mean value of the corresponding 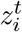 under the Wright-Fisher model with selection conditioned in the same way. Note that the Wright-Fisher model is a special case of a Cannings model with *σ*^2^ = 1. By imposing that the two processes correspond at the level of the means, we are ensuring that the only difference between the Wright-Fisher model and the Cannings extension is the degree of random fluctuations.

Lastly, we need a sampling procedure to obtain a set of coalescence times from these models. To do this, we can consider the coalescence times of a finite random sample of *k* selectively advantageous individuals at the final time-point. The only way to guarantee we are able to sample a set of *k* selectively advantageous individuals is to condition the mutation on having expanded to at least *k* individuals at the final time-point, which is equivalent to conditioning on fixation when the total number of generations *L* is large. Throughout the paper, we will express this assumption by conditioning on eventual fixation. Although we assume eventual fixation, we are interested in cases where the *k* individuals are sampled during the selective sweep or shortly after fixation. Importantly, the assumption that we have sampled from the class of selectively advantageous individuals is not necessary, and we could have assumed that the *k* individuals are randomly sampled from the entire pool of individuals and our results will not change.

We can now state the question more precisely. Suppose we have observed a set of coalescence times generated by some Cannings model with selection that we defined above. Now suppose we infer a posterior distribution for the selection coefficient and time of onset of the selectively advantageous mutation from the observed set of coalescence times using a different Cannings model with an arbitrary *σ*^2^. What will happen to the inferred distributions if we change *σ*^2^ in the model used for the inference?

We can take a step back and ask what coalescent theory would predict. In particular, we can infer the effective population sizes from an observed set of coalescence times [9, 10, 11, 12]. We can then consider the robustness of inferences of *s*. To begin, we will consider the Wright-Fisher model with selection that we defined (rather than an arbitrary Cannings model for simplicity, but the concepts will still hold), and then define *n*(*t*) as the number of individuals with a selectively advantageous mutation as a function of time in units of generations. Note that *n*(*t*) is a random variable, which produces a well-defined coalescent rate 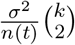 [10, 11, 12], where *k* is the number of sampled individuals, after *n*(*t*) is realized. We can consider the inferred effective population size from a set of coalescence times at two different generations 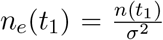 and 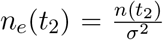, where *n*_*e*_(*t*) is the effective population size as a function of time. The selective coefficient can be indirectly inferred from the estimator 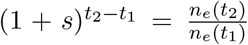 which does not depend on the reproductive variance. Since *σ*^2^ is not required to infer *s*, then naively we can argue that inference of *s* is robust. However, this argument does not say anything about the fluctuations in the inferred value of *s*. The inference results in a posterior distribution of *s*, not a single value. It is conceivable that the larger the assumed *σ*^2^ in the model, then the larger the uncertainty in 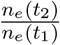. However, we will show later that indeed the inferred posterior distribution of *s* is independent of *σ*^2^.

We can also consider the robustness of the inferred time at which the selectively advantageous mutation arose. We can begin by considering the inferred population size trajectory which is *n*(*t*) = *σ*^2^*n*_*e*_(*t*), where *σ*^2^ is the reproductive variance of the model used for the inference. When the Wright-Fisher model with selection is conditioned on fixation, as a crude approximation, the selectively advantageous mutation grows according to 1 + *t* on average until a population size of 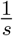 is reached, at which point selection is stronger than drift and the mutation becomes destined for fixation. This is known from modeling average frequencies of mutations using birth-death processes [56, 57], as well as the probability of fixation [58, 59], and considering the scale at which different terms dominate, such as in [38]. Using coalescent theory, we will infer a population size trajectory of *n*(*t*) ≈ *σ*^2^(1 + *t*) for small t values, on average, from an observed set of coalescence times generated from the Wright-Fisher model. Importantly, when *σ*^2^ ≠ 1 in the model used for inference, the inferred population size reaches one at a different generation: by setting *σ*^2^(1 + *t*) = 1 we derive 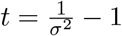 which is a number in between −1 and 0 provided *σ*^2^ ≥ 1. This is different from the time at which the true population size of the Wright-Fisher model (where *σ*^2^ = 1) reaches one on average, which is at generation *t* = 0 by definition, and differs from the time at which the inferred population size is one by at most the duration of one generation when *σ*^2^ ≥ 1, but can be larger than one generation if *σ*^2^ *<* 1. Again, we run into the same problem that arises in the inferred selection coefficient: this naive argument neglects the fluctuations in the inferred time of onset. Increasing *σ*^2^ increases the random fluctuations in the model which increases the uncertainty in the inferred time. We will later derive explicit formulas describing the offset of the inferred posterior under different *σ*^2^ that fully accounts for these random fluctuations.

Up to this point, we have only considered the impact of varying *σ*^2^ in the model, but have not considered the impact of varying the assumed *L* on the inferred posterior of *s* and *g*. To better understand the impact, it is most instructive to convert the branch lengths of the data tree to an absolute time-scale (such as units of minutes, days, years). This is oftentimes done empirically if the branch lengths are in number of mutations and we know the mutation rate per unit time. The model assumes an arbitrary number of generations that spans the tree *L*, but the absolute amount of time spanning the tree is assumed to be a known, fixed quantity *a*. If 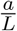 is the duration of a generation in the model, then the coalescent rate in units of absolute time is 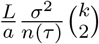. Note that we can define 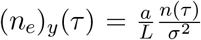 as the effective population size with a *σ*^2^ converted from variance per generation to per unit time, since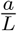 is a conversion factor. For simplicity, we can write 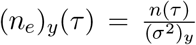, where 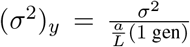, and will use the subscript *y* to denote parameters converted to units of an absolute time-scale throughout the paper. We can then define an estimator for the inferred selection coefficient in units of time 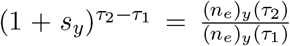 and note that the same arguments apply. In particular, the inferred *s*_*y*_ is independent of (*σ*^2^)_*y*_, and consequently independent of *L*. Similarly, the inferred population size during drift of a tree generated from a Wright-Fisher model where the duration of a generation is assumed to be one is *n*(*τ*) = (*σ*^2^)_*y*_ (1 + *τ*). Note that it follows that the inferred time is 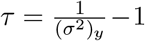, which shows an impact of at most the duration of a generation for (*σ*^2^)_*y*_ ≥ 1, but can be larger than one generation whenever (*σ*^2^)_*y*_ *<* 1. Again, these arguments ignore the impact of random fluctuations due to a different (*σ*^2^)_*y*_ (or equivalently *L* and *σ*^2^ in the model). Similar to before, we will derive explicit formulas showing the offset of the inferred posterior under different *L* or *σ*^2^ that fully accounts for the random fluctuations.

The reader can verify that these arguments hold for varying *L* only when the inferred parameters have units of an absolute time-scale, and not when they have units of generations. For the remainder of the paper, we will derive the results on an absolute time-scale to be as general as possible. This is because any formula expressed in units of absolute time can be converted to units of generations by assuming 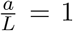. To derive explicit formulas, we will need to take a more rigorous approach and model the fluctuations in gene-frequencies as a function of time. This requires invoking the Kolmgorov description of the Cannings model class.

## 4 Scaling limit of the Kolmogorov forward equation beginning from a population size of one

In the previous section, we presented arguments for the robustness of the inferred *s*_*y*_, *g*_*y*_ to the assumed *σ*^2^ or *L* in the model. These arguments failed to account for the contributions of random fluctuations of the model to the posterior distribution. To account for random fluctuations, we will use the Kolmogorov forward equation [6] to model the changes in gene frequencies through time. This approach is a diffusion approximation, first used by [60, 1] as well as in various influential works more explicitly, including [61, 62, 63, 64] and is currently a standard approach in population genetics. We will consider a particular limit where the error in the posterior of *s*_*y*_, *g*_*y*_ under different *σ*^2^ or *L* completely vanishes and use it as an example to build intuition. This will give us a deeper understanding of the problem, help us identify where robustness breaks down, and point us towards an approach to carry out rigorous calculations.

We can begin by considering the Kolmogorov forward equation for the Cannings model. Here *φ* represents the probability density of gene frequency *x* at generation *t*, where the remaining parameters have the same definitions as in the Cannings model defined. In particular, the parameters of the equation are *N*, the total population size, *σ*^2^, the variance in offspring number, and *s*, the selection coefficient. The fourth parameter is *g*, the number of generations that has elapsed from the initial time-point when the clone size is one to the final time-point. *g* is implicitly incorporated as the duration of time for which we run the forward Kolmogorov equation. Namely, *φ*(*x*, 0) = *δ*(*x*− 1*/N*), and *k* individuals are sampled at the final time-point *g* from a realization of the clone whose frequency is described by *φ*(*x, t*). As before, our goal is to infer *s* and *g* from the coalescent structure of the *k* individuals. The dynamics of *φ*(*x, t*) is given by,

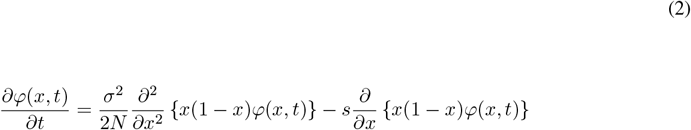

As suggested before, we will convert the equation to units of absolute time. This can be done using the differential 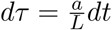 and the chain rule to obtain

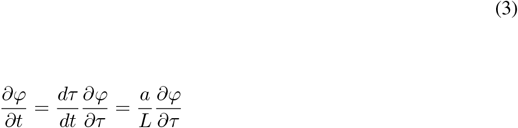

and substitute (Eq. 3) into (Eq. 2) and rewrite as follows

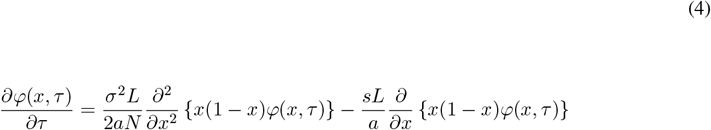

We can write the above equation in terms of *s*_*y*_, (*N*_*e*_)_*y*_. This can be done by first noting that if there are 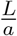 generations per unit time, and the population grows by a factor of 1 + *s* each generation, then the population growth is 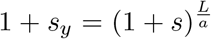 per unit time. In addition, we can use 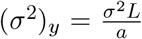, and write *N* as an effective population size 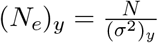. Substituting these values and assuming |*s*| *<<* 1 we obtain:

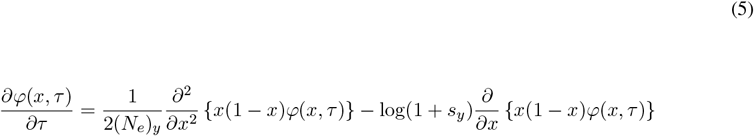

It is clear that this expression only depends on *L* and *σ*^2^ through (*N*_*e*_)_*y*_ and *s*_*y*_. Therefore, we will obtain the same solution if we arbitrarily change *L* and *σ*^2^ while keeping *s*_*y*_ and (*N*_*e*_)_*y*_ constant. For example, if we let *L* → *cL*, then 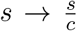 so that *s*_*y*_ remains constant, and *N* → *cN* so that (*N*_*e*_)_*y*_ remains constant. Similarly for *σ*^2^ → *cσ*^2^, *N* → *cN* so that (*N*_*e*_)_*y*_ remains constant, and *s*_*y*_ remains constant merely by the fact that it does not depend on *σ*^2^.

In addition, it follows that the distribution of coalescence times of a random sample of selectively advantageous individuals is also preserved as long as *s*_*y*_ and (*N*_*e*_)_*y*_ are held fixed. For example, we can define the random variable *x*(*τ*) as the frequency (i.e. fraction of *N*) of selectively advantageous individuals as a function of time described by (Eq. 5). The effective population size of the fit subpopulation is then *x*(*τ*)(*N*_*e*_)_*y*_, and the rate of coalescence at each time-point is 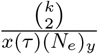 after any realization of *x*(*τ*). Since the distribution of *x*(*τ*) and thereby 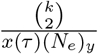 is preserved, then the distribution of coalescence times is also preserved when changing *L* or *σ*^2^ as long as *s*_*y*_, *g*_*y*_, (*N*_*e*_)_*y*_ are held fixed, where *g*_*y*_ is the duration of time in absolute time units we run *x*(*τ*) under (Eq. 5) before sampling.

Although this scaling is well-understood, it is useful to recall its biological meaning. In particular, increasing *L* → *cL* or *σ*^2^ → *cσ*^2^ increases the rate of diffusion of gene frequencies by increasing (*σ*^2^)_*y*_. However, the scaling also requires *N* → *cN* to preserve (*N*_*e*_)_*y*_, which decreases the rate of diffusion since the rate also depends on absolute population size, thereby canceling the increase to (*σ*^2^)_*y*_. In addition, changing *L* → *cL* requires 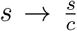 to preserve *s*_*y*_, so the increase in the growth rate due to adding more generations is balanced by a decrease in selection at each generation. Put more simply, under this scaling we are preserving the balance of drift and selection on an absolute time-scale by requiring *N* and *s* to cancel any change in *σ*^2^ or *L*. This scaling also ensures that the effective population size of the selectively advantageous subpopulation *x*(*τ*)(*N*_*e*_)_*y*_, and hence the coalescence rate, is preserved.

Up until now, we have pointed out a standard scaling of the Kolmogorov forward equation, and how it coincides with the Kingman coalescent. It may be tempting at this point to assume that we have solved the problem. For example, we could naively argue that since any combination of the parameter values *s*_*y*_, *g*_*y*_, (*N*_*e*_)_*y*_ produces the same distribution of coalescence times, then it follows that we can infer these parameter values independent from the choice of *L* or *σ*^2^. Another way of saying this is that the posterior distribution will not change if we change the *L* or *σ*^2^ in the model used for the inference. However, this argument is actually incorrect. In particular, the Kolmogorov forward equation is expressed in terms of a frequency *x*, and as a result, the initial condition is also expressed as a frequency. More precisely, the initial condition can be written as

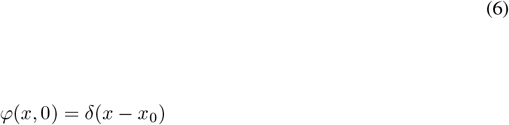

where the right hand side of (Eq. 6) is a Dirac delta function, and *x*_0_ is the starting frequency. However, our model does not assume a fixed initial frequency. It instead assumes an initial frequency of 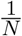, since the process begins from a fixed population size of one. This implies that the initial condition changes with the scaling we considered above. For example, we can rewrite our initial frequency in terms of the parameters of the equation in absolute time units and *σ*^2^ and *L* to obtain 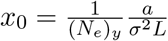, and hence

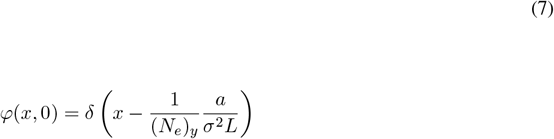

It follows that the initial condition changes with *σ*^2^ and *L* while holding fixed *s*_*y*_, *g*_*y*_, (*N*_*e*_)_*y*_ although the equation itself is invariant, which implies the solution of the Kolmogorov forward equation (i.e. the distribution of gene frequencies or *x*(*τ*)) changes under the scaling. As a result, such an invariance does not actually hold under these assumptions.

We now consider the invariance of the Kolmogorov forward equation with the correct initial condition of one. We will show that the invariance under changing *L* and *σ*^2^ while keeping fixed (*N*_*e*_)_*y*_ and *s*_*y*_ still holds when conditioned on eventual fixation in the limit *L* → ∞ or *σ*^2^ → ∞. We will use this result as a starting point to think about what happens when *L* and *σ*^2^ are finite.

To begin, denote the density conditional on eventual fixation as *φ*(*x* | *F*). Using Bayes’ theorem, we can write

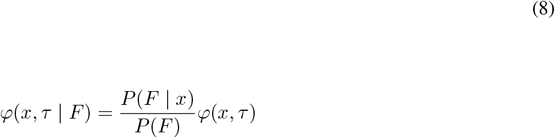

where we have set *P* (*F* | *x, τ*) = *P* (*F* | *x*), since the probability of eventual fixation is time-independent. The probability of fixation of a mutation with frequency *x* within a sufficiently large population of size *N* and with selection coefficient *s* is given by [59]. Therefore, after rewriting the parameters in terms of *s*_*y*_, (*N*_*e*_)_*y*_

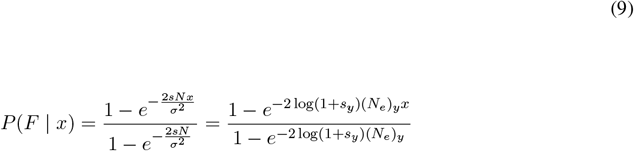

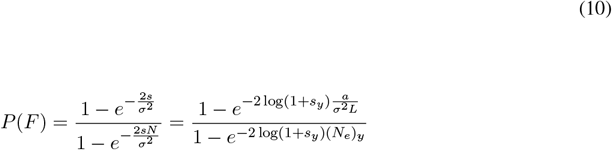

Note that (Eq. 10) is (Eq. 9) with 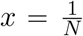, since we always begin with one selectively advantageous individual. Substituting (Eq. 9-10) back into (Eq. 8) gives us

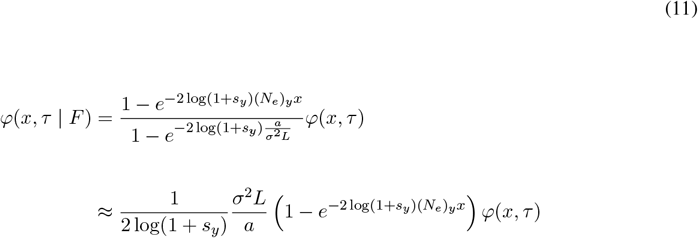

where we have assumed that 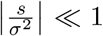. This limit assumes that drift is stronger than selection initially when the population size is small. The unconditional gene frequency density *φ* is given as the solution to the diffusion approximation [7], which in our notation is

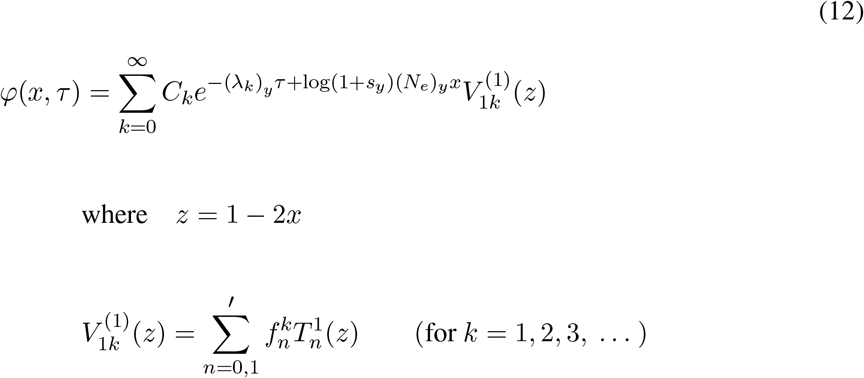

The primed summation is over even values of *n* whenever *k* is even, and odd values of *n* whenever *k* is odd. 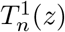 are Gegenbauer polynomials of the first kind [65] and are defined as

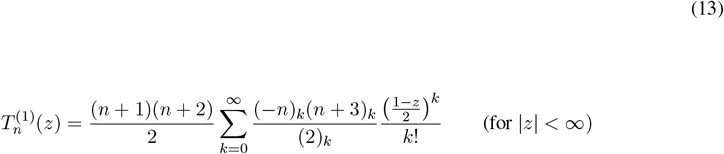

where we have used Pochhammer notation to denote a rising factorial defined as

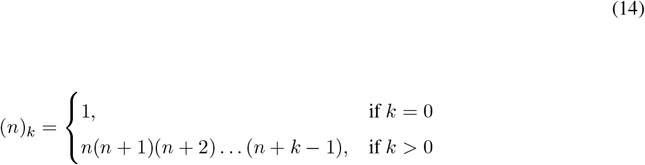

The (*λ*_*k*_)_*y*_ are eigenvalues, and 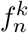 are constants which depend on *s*_*y*_ and (*N*_*e*_)_*y*_ and *k*. The eigenvalues and constants arise through a related oblate spheroidal angle equation [66, 67, 68]. These values can be numerically approximated using the method of continued fractions or taken from tabulated values [68, 69].

The *C*_*k*_ are determined by the initial condition *δ*(*x* − *x*_0_) and are given by

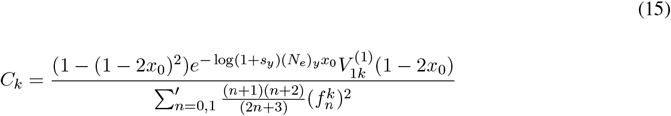

We are interested in the initial condition 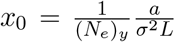. We can substitute the initial condition into (Eq. 15) and rewrite it in a way that will allow us to ignore second order terms

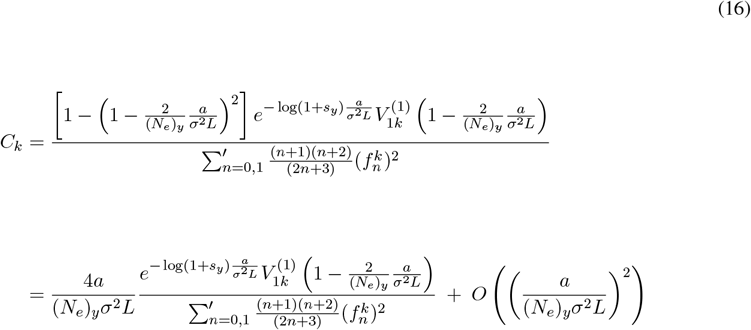

By substituting (Eq. 16) into (Eq. 12) and distributing the sum we obtain

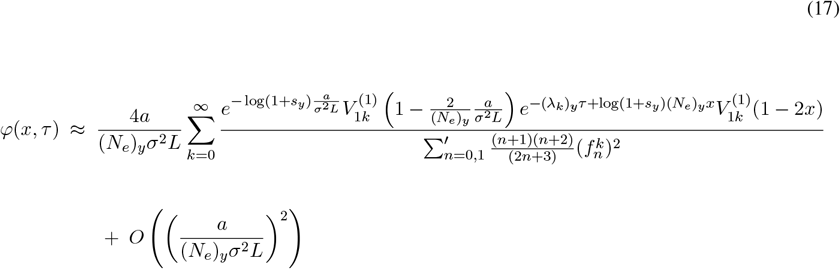

Lastly, we can substitute (Eq. 17) into (Eq. 11) and obtain

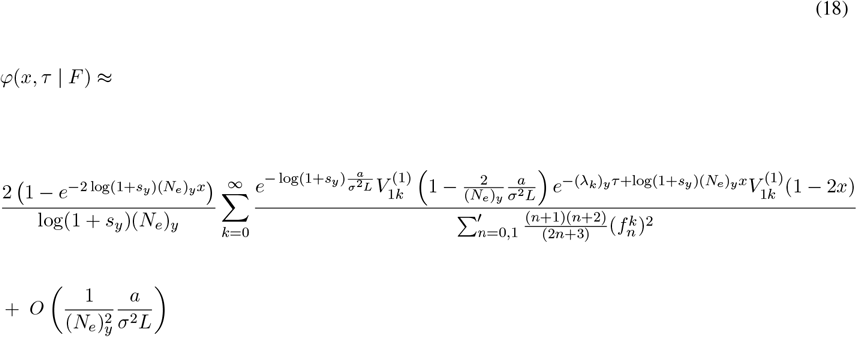

Note that in this form, we can see that the sum converges as *L* → ∞ or *σ*^2^ → ∞ to a limit function of *x, τ, s*_*y*_, (*N*_*e*_)_*y*_ that is independent of the initial condition 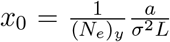. This suggests that the process forgets the initial differences in frequency at *τ* = 0, while preserving the distribution of frequencies soon after. As before, this implies that the effective population size *x*(*t*)(*N*_*e*_)_*y*_ of the selectively advantageous subpopulation is preserved, and hence the distribution of coalescence times is also preserved, showing robustness to changes in the initial frequency caused by changes in *L* and *σ*^2^. Another way of saying this is that the posterior distribution converges to the following limit function

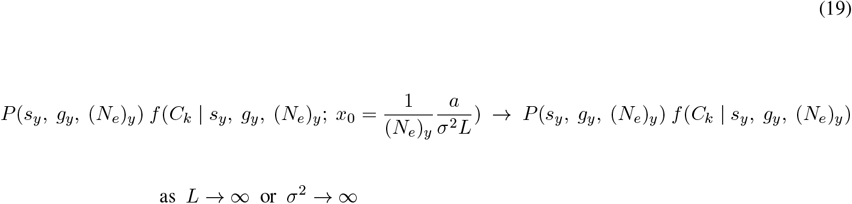

where *P* (*s*_*y*_, *g*_*y*_, (*N*_*e*_)_*y*_) is a prior distribution, 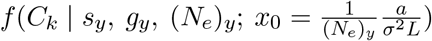 is the likelihood function, *C*_*k*_ are a set of observed coalescence times, and we have ignored the constant of integration or can think of it as being absorbed by *f*. The likelihood function on the right hand side is denoted as being independent of the initial condition, and as a result so is the constant of integration.

Such a result may not be too surprising upon further reflection. We can think about the particular scaling more carefully and begin by first writing the fitness in absolute time units as

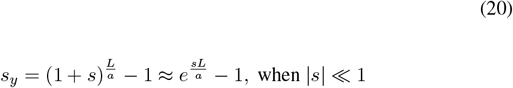

If we let *L* → *cL*, then we must have 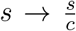to keep *s*_*y*_ constant. However, conditional on fixation, the population size required for escaping extinction during drift (i.e. becoming established for fixation due to selection) goes from 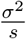to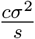. Both processes take on average 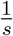 generations to establish, or in absolute time units a duration of 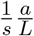, which remains constant under the scaling [56, 57, 58, 59, 38]. This implies the population size increases on average by a factor of *c*. However, *N* → *cN* to keep (*N*_*e*_)_*y*_ constant. Since both the mutant population size and the total population size both increase by a factor of *c*, the frequency is preserved under this scaling. As we argued before, since *x*(*t*) and (*N*_*e*_)_*y*_ is preserved, the effective population size *x*(*t*)(*N*_*e*_)_*y*_ and hence the distribution of coalescence times is preserved. A similar argument can be made for *σ*^2^ → *cσ*^2^. Under this scaling, the population size at which the population is established is 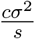 while the time to establish is independent of *σ*^2^ and constant. Since *N* → *cN* to keep (*N*_*e*_)_*y*_ constant, *x*(*t*) is preserved, and it follows that the distribution of coalescence times is also preserved.

These dynamics are, of course, well understood. However, it’s worth emphasizing that there is a key distinction between the original scaling we considered which assumes a fixed initial frequency vs an initial population size of one. In the first case, the scaling results in the mutant population size increasing by a factor of *c*, but only because the initial population size is increased by a factor of *c* by definition. In the second case, the population size is scaled by a factor of *c* because the scaling results in an increased rate of extinction which means the trajectories that are observed are the ones that fluctuated to a larger population size and quickly escaped drift early on. In other words, the rescaling of population size is related to the dynamics of the population, and not a result of arbitrarily changing the initial population size. Importantly, this rescaling of the mutant population size has nothing to do with the fact that *N* → *cN*, and only related to *s, L* and *σ*^2^ which determine the time and population size at which drift is escaped. The reason *N* → *cN* is required in the Kolmogorov equation is to preserve the time at which saturation is reached, but the scaling of trajectories is fundamentally a result of the underlying dynamics of the selectively advantageous subpopulation.

This invariance does not just apply to coalescent tree data. We have already established that the distribution of *x*(*t*) is invariant, so the results would apply if our data were a set of gene-frequencies *x*(*t*) measured at time-points after the initial advantageous mutation arose. Similarly, the observed site-frequency spectrum of the selectively advantageous subpopulation is also invariant. This is because the combined factor of increase in population size over time and the equivalent factor of decrease in the probability of fixation implies that the same number of neutral mutations that arise at a given time-point and reach the final time-point is observed in both processes (any process and the scaled parameter version of it). Extensions to these other data forms should ultimately be calculated and studied more rigorously. We will continue to focus on coalescent trees since our goal is not to create a catalog of all possible forms of data under which these results apply, but to present a framework/procedure that can be easily adapted to other forms of data beyond coalescent trees.

As we previously argued, the invariance under an initial population size of one is only true in the limit of *L* → ∞ or *σ*^2^ → ∞. In this limit, the scaled process when conditioned on fixation drifts to the exact initial frequency of the unscaled process with vanishing fluctuations so that the two processes are described by the same solution to the Kolmogorov forward equation. For finite *L* and *σ*^2^, coalescence events that occur near the time at which the selectively advantageous mutation arose will have subtle differences in their distribution of coalescence times under different values of *L* and *σ*^2^. As a result, the coalescence times will retain their invariance only after some *τ* = *ϵ*, where *ϵ* → 0 in the limit as *L* → ∞ or *σ*^2^ → ∞. Although it is possible to consider the rate of convergence of *ϵ* directly, it is not a particularly useful quantity, since we are interested in explicitly calculating how the posterior distribution is impacted by this initial error. From this approach, it is neither clear what the impact should be nor does it even suggest a way to compute it. What we can say though is that the impact could in theory be significant. For example, we can consider the case of the inferred times at which blood cancer mutations arise in individual patients [33, 34] where the time between generations is thought to be one year [70]. If, for example, the error happens to be a fixed number of ten generations, then the inferred age of onset of the mutation would be impacted by ten years which could significantly change the interpretation of the results. Such an error would, of course, vanish as the number of generations per unit time becomes infinite. To directly compute the impact on the posterior distribution, we will need to introduce new concepts. In particular, we will need to map ensembles of trajectories from different models with different initial conditions in a way that preserves coalescence statistics. To do so, we will solve a first passage time problem.

## 5 Robustness is a first passage time problem

So far we have shown that under the appropriate scaling of parameter values and under certain limits, namely where *L* → ∞ or *σ*^2^ → ∞ such that (*N*_*e*_)_*y*_ and *s*_*y*_ are held constant, then the posterior distribution converges to the following limit function

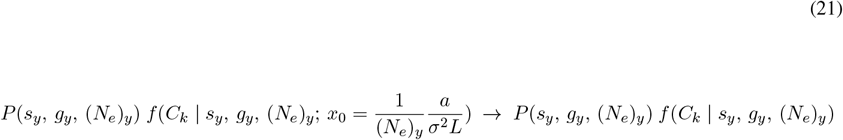

which is independent of the initial condition. Recall that *g*_*y*_ is the time of onset of the clone or the absolute time from the occurrence of the selectively advantageous mutation (when the clone arises with population size one) to the time at which random individuals are sampled from the clone to construct their coalescent tree. Although the posterior becomes arbitrarily close in the limit, it will differ from the limit function for finite *σ*^2^ or *L*. To compute the impact, we will first discuss how to correctly establish a correspondence between trajectories across different scalings, and in doing so it will become clearer how to compute the error.

We will consider again the first scaling in this paper, which assumed a fixed initial frequency rather than a fixed initial population size of one. We will consider two processes that differ by the scaling *L* → *cL*. We will assume that the processes have an initial population size of 1 and *c* (*c >* 1) for the first and second process respectively to ensure the frequency remains fixed as *N* → *cN*. We will refer to the first process as the original or unscaled process and the second as the scaled process. The invariance of the Kolmgorov forward equation suggests the following: for any trajectory in the original process, there exists an equivalent trajectory in the scaled process that is precisely a factor of *c* larger in population size at every time-point and has an identical probability of occurring. Since effective population size is preserved across both trajectories, the distribution of coalescence times from a random sample of either trajectory is the same.

Now consider two such pairs of trajectories (an original and its scaled counterpart). Since the scaled trajectory begins from a population size of *c* rather than 1, we can use it to create a new trajectory that begins from a population size of 1 by stitching it together with another trajectory that began at a previous time point, but reaches a population size of *c* for the first time and terminates precisely at the time at which the scaled trajectory begins. We stress that it is important to only consider trajectories that reach *c* for the first time, thereby excluding those that have become or exceeded a population size of *c* at any time-point before. Importantly, we can create an ensemble of trajectories out of the scaled trajectory, each with a different probability of occurring. For example, a trajectory that begins much farther back than the scaled trajectory is unlikely to reach *c* for the first time when the scaled trajectory begins, since reaching *c* before is likelier. By design, each trajectory of the scaled ensemble has an identical probability of any *C*_*k*_ that the original/unscaled process can produce, since the effective population size matches the unscaled process after the time-point at which it begins.

These ideas can be restated in terms of frequencies. In particular, the unscaled process has a starting frequency of 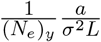 while the scaled ensembles begin farther back from a starting frequency of 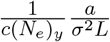and reach 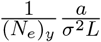 precisely when the unscaled process begins. Conditional on reaching 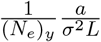 when the unscaled process begins, both processes will have the same trajectories of frequency from there on and hence the same distribution of coalescence times. For the remainder of the paper, we will use the language of frequencies because it simplifies the notation.

Establishing this correspondence allows us to associate the trajectories of any process with an ensemble of trajectories from a scaled version of the process in a way that preserves coalescence statistics. This is useful because we can generate a posterior distribution of trajectories using a model with scaled parameter values with respect to an underlying process, and map them to trajectories from a posterior distribution generated using the correct parameter values. The mapped trajectories across the posteriors will only differ in that the ensembles from the scaled posterior will have begun farther back, which means that the impact of the scaling on the inferred *g*_*y*_ is related to the time it takes the scaled process beginning from a smaller frequency to reach the starting frequency of the original process. This is known as a first passage time problem which is a recurrent theme in stochastic processes.

Importantly, this offset will have no impact on the inferred posterior of fitness *s*_*y*_. This is because the time it takes the scaled process to reach the frequency of the original process is independent of its fitness *s*_*y*_, since this occurs during the drift phase long before selection determines the dynamics of the trajectories. By the time *s*_*y*_ impacts the dynamics of the trajectories, the scaled trajectory has reached the frequency corresponding to the start of the unscaled trajectory. Drift switches over to exponential growth at a population size of 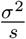. We live in the regime 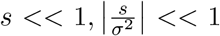 where these assumptions are clearly satisfied, implying that the scaled trajectories match the starting frequency of the unscaled trajectory long before drift ends. Therefore, the posterior of *s*_*y*_ will not change if *L* and *σ*^2^ change as long as (*N*_*e*_)_*y*_ and *s*_*y*_ are held constant.

We will now make these intuitions rigorous and use it to come up with a more sophisticated definition of robustness. Our calculations will focus on the posterior of *g*_*y*_ which requires an explicit calculation. The invariance of the inferred *s*_*y*_ follows as a direct consequence of the way we have associated trajectories across the different processes.

## 6 Definition of Robustness

We will now express the ideas in the previous section in rigorous mathematical terms. We will derive a set of equations that describe the impact on the posterior distribution under changing model assumptions *σ*^2^, *L* and take them as our definition of robustness. We will then derive under certain conditions a set of simple expressions relating robustness to the mean and variance of the time of first passage. As we will see, this approach is both conceptually useful and will make the assumptions more explicit.

We will begin by deriving a posterior distributions using a (*σ*^2^)_*y*_ assumed to match the true underlying process, and compare it to a posterior distribution using *c*(*σ*^2^)_*y*_. Note that the scaling (*σ*^2^)_*y*_ → *c*(*σ*^2^)_*y*_ is equivalent to either *σ*^2^ → *cσ*^2^ or *L* → *cL*, but we will now use the former notation as it encompasses both. The changing initial condition will now instead be expressed as 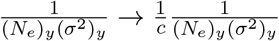. We can consider the marginal posterior distribution of the inferred time at which the selectively advantageous mutation arose (i.e. time at which the population was originally one) for the unscaled and scaled process which we will denote as 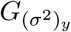 and 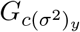. More precisely 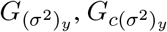 are random variables which denote a sampled value from these posteriors. We are interested in deriving an expression for 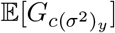 in terms of 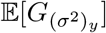, and 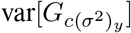 in terms of 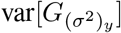, although generalizing to higher moments is possible.

The distribution of 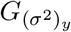 we will denote by 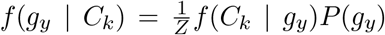, where *P* (*g*_*y*_) is the prior distribution of the time of onset, and 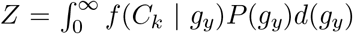 is the integration constant. For the scaled process, we will denote the distribution of 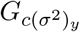 as 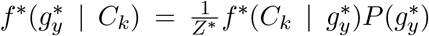, where 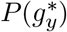 also denotes the prior distribution with respect to 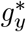, but have assumed the prior distributions in both processes are the same, namely that 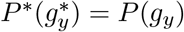 whenever 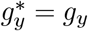, and that 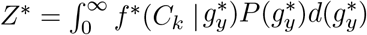. Note that 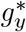 and *g*_*y*_ both refer to the time (in absolute time units) into the past at which the population was initially one in both processes. Next we will denote the probability density that the scaled process with parameter *c*(*σ*^2^)_*y*_ reaches frequency 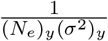 for the first time after a duration of time *T* beginning from frequency 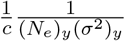 as *Q*(*T*). Importantly, as we stated in the previous section, the distribution of coalescence times will be the same across both processes if we condition on the frequency of the scaled process reaching 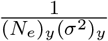 when the original process begins. If we denote the trajectory of frequency of the scaled process as *x*^*^ where 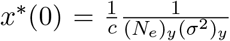, then we can denote this assumption by conditioning the likelihood function and writing: 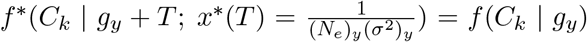 for all *T >* 0. This assumption will allow us to express 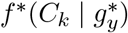 in terms of *f* (*C*_*k*_ | *g*_*y*_) and *Q*(*T*).

Before doing the calculation, we will take a quick digression to note that this assumption can break down. For example, if the original process begins after the final coalescent event (here, after means closer to the present), but the scaled process begins before, then the original process will have a probability of zero of producing the observed tree and the scaled process will have a nonzero probability, yet we will have assumed that both processes have equal probability of producing the observed tree. However, this assumption appears not to significantly impact the results as simulations will show in a later section. This is likely because coalescence events rarely occur in the time-interval before the population size reaches *c*. In addition, we will end up integrating the likelihood of coalescence across different times of onset, and most of the contribution to the integral will occur at time-points before the final coalescence event which is when the assumption does not break down.

We can now write down the posterior of 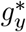, and use the law of total probability along with this assumption to obtain:

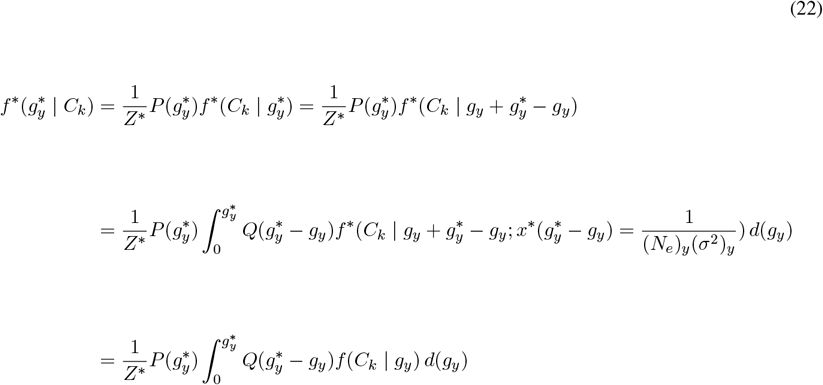

Here we have employed the use of an integral to simplify the notation and avoid having to sum over fractional durations of time. Finally, note that the integration constant is

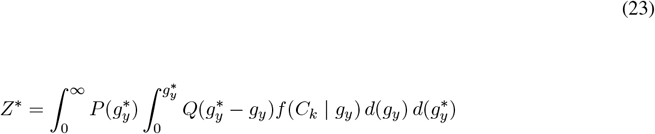

We can use (Eqs. 22-23) to compute the expectation and variance. We will begin with the expectation:

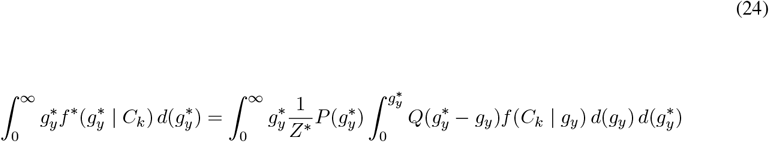

Switching the order of integration we obtain:

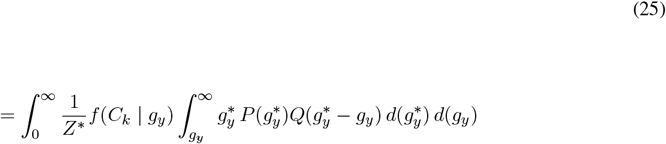

Next we do a change of variables 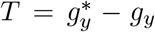 in the inner integral, which can be done since *g*_*y*_ is constant:

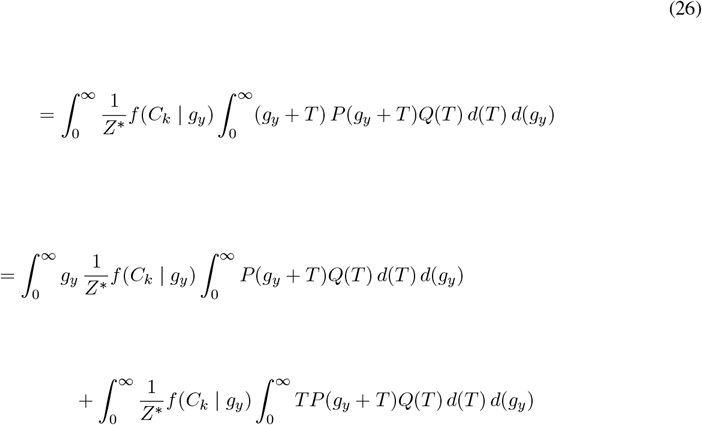

We can also rewrite the integration constant in a similar manner by changing the order of integration and using the same change of variables 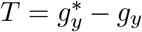:

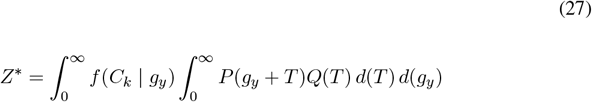

If we assume that *P* (*g*_*y*_ + *T*) = *P* (*g*_*y*_), or equivalently that it is a uniform distribution, then from (27) we see that *Z*^*^ = *Z*, and from (26) we obtain

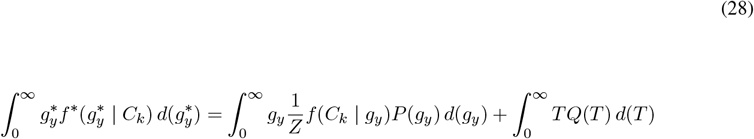

or equivalently

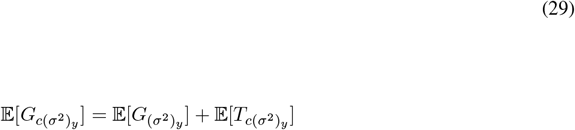

where 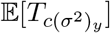 is the expectation of the first passage time to 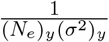 using the scaled process and with starting frequency 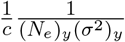

We can then compute the second moment of 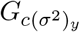 in a similar fashion, again with the assumption *P* (*g*_*y*_ + *T*) = *P* (*g*_*y*_), and use it together with the computed mean to obtain the variance:

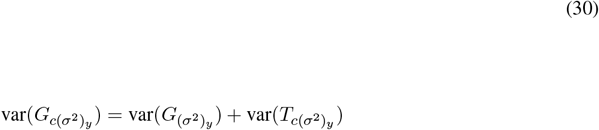

We will not compute the higher moments although it is possible to continue this procedure.

The equations above show the relationship between robustness and the time of first passage. Furthermore, under the simplifying assumption of a uniform prior distribution, 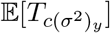 and 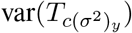 are exactly the offset of the mean and variance of the posterior distribution respectively. Therefore, in this case, the problem of robustness reduces to finding the mean and variance of the time of first passage. We will consider non-uniform prior distributions in a later section. For now, we will compute the offset of the mean and variance assuming *P* (*g*_*y*_ + *T*) = *P* (*g*_*y*_).

It is important to note that these equations hold for only *c* ≥ 1, since we are implicitly assuming the process with a larger (*σ*^2^)_*y*_ and hence smaller starting frequency is begun farther back (in this case the process with parameter *c*(*σ*^2^)_*y*_). However, we can generalize the equations by considering the case 0 *< c <* 1. The key insight is to notice that these equation more generally show how inferred values differ between two inferences with different assumed values of (*σ*^2^)_*y*_, independent of whether either (*σ*^2^)_*y*_ actually matches the variance in the underlying process. We considered the unscaled (*σ*^2^)_*y*_ as the true underlying process in our derivations, but there is nothing inherent about this in the equations. More generally, the left side of the equation corresponds to the inference with a larger (*σ*^2^)_*y*_ while the left term of the right hand side corresponds to an inference with a smaller (*σ*^2^)_*y*_. This allows us to write for 0 *< c <* 1:

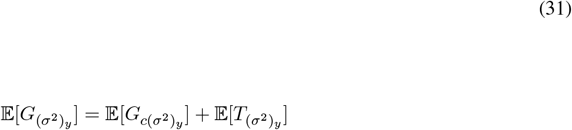

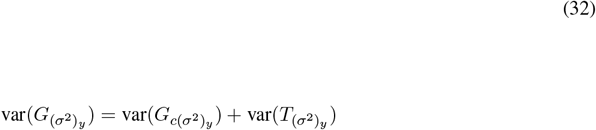

Or equivalently

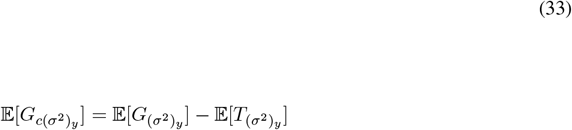

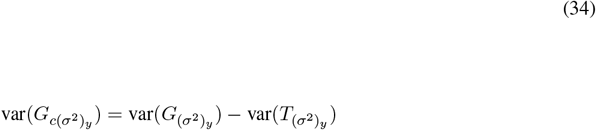

We can now interpret (*σ*^2^)_*y*_ as the correct parameter of the underlying process, and obtain an expression for the impact of using a model with 0 *< c <* 1. Apart from the obvious difference that the correction is negative in sign, the first passage correction is now computed using the parameter value of the underlying process (*σ*^2^)_*y*_ and the time it takes to reach a frequency of 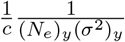 beginning from a frequency of 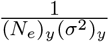.

## 7 Calculating the time of first passage

In the previous section, we derived a set of equations to describe the offset of the mean and variance of the posterior of *g*_*y*_ from assuming an incorrect (*σ*^2^)_*y*_ in the model. Under the assumption of a uniform prior distribution, we showed that the offset is the expectation and variance of the time of first passage. We will now attempt to compute these quantities directly. In particular, we will compute 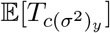 and var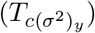 for *c* ≥ 1 and 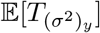 and var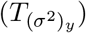 for 0 *< c <* 1, where 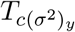 is the first passage time distribution from 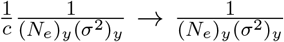 using the model parameter *c*(*σ*^2^)_*y*_, and 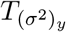 is the first passage time distribution from 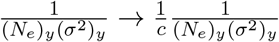 using the underlying process parameter (*σ*^2^)_*y*_. To derive formulas that hold for both cases, we will denote the initial frequency of the process that is pushed back as *p*, the frequency that must be reached as *f*, and substitute the correct values later on. To calculate the mean and variance of the time of first passage, we will follow an approach that was used to compute the moments of the time until fixation by directly integrating the Kolmogorov backwards equation [71, 6], but with modified conditions. However, other approaches are possible that consider the same question [1, 63, 59, 72, 73, 74, 75, 76, 77, 78, 79, 80]. We will begin by writing down the Kolmogorov backwards equation for the Cannings model [6, 59]. As an approximation, we will drop the selection term that depends on *s*_*y*_ since the first passage occurs during the drift phase before the mutant population is established.

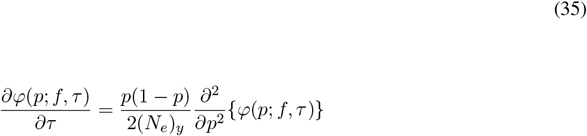

Here *φ*(*p*; *f, τ*) represents the probability that the frequency of the mutant subpopulation takes on a value of *f* (0 *< p < f* ≤ 1), where *f* is an absorbing boundary, at generation *τ* beginning from a frequency of *p*. The most widely studied case is *f* = 1, where *φ*(*p*; 1, *τ*) represents the probability of an allele having reached fixation [59, 71, 72]. The actual Cannings trajectories do not remain at frequency *f* once *f* is reached for 0 *< f <* 1, but artificially introducing an absorbing boundary allows the process to retain information on the fraction of trajectories that reach *f* for the first time at any time-point. Importantly, the equation above does not explicitly depend on *f* because the absorbing boundary is specified in the boundary conditions, even in the case of *f* = 1. In particular, we must assume:

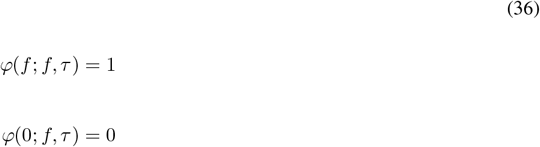

For simplicity of notation, we will now write *φ*(*p, τ*), where it is implicit we are modeling the probability of reaching *f*.

We are interested in computing the mean and variance of the distribution of time it takes to reach *f* for the first time beginning from *p*, conditioned on the process reaching *f* eventually. The reason we exclude the trajectories that go extinct before reaching *f* is because it is only the trajectories under the scaled parameter values that reach a frequency *f* that will have the same coalescence time distribution as the original process. Recall that we have chosen to model *f* as an absorbing boundary. The precise justification for modeling *f* this way is as follows: it allows us to first compute the probability that a Cannings trajectory reaches *f* at some point (rather than going extinct before reaching *f*) through lim_*τ*→∞_ *φ*(*p, τ*) = *φ*(*p*), and then to compute the probability of reaching *f* by time *τ* conditional on the process eventually reaching *f*, 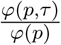. This quantity 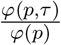 is important because it is the CDF of the first passage time conditional on *f* being eventually reached. This follows from the definition of an absorbing boundary, since any trajectories that reach *f* at any given time are by definition doing so for the first time. The probability density of the first passage time is then 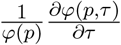. We will first focus on computing *φ*(*p*) and then compute the mean and variance of the density. To compute *φ*(*p*), we will take the limit as *τ* → ∞ of the Kolmogorov backwards equation so that 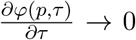, which follows from the fact that 0 or *f* is eventually reached. We therefore obtain

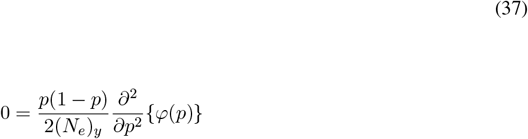

which implies

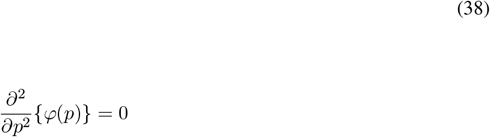

We then integrate and apply the boundary conditions *φ*(*f*) = 1, and *φ*(0) = 0 to obtain

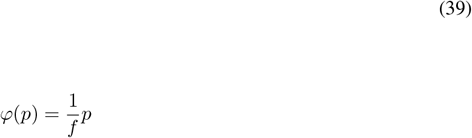

Note that when *f* = 1 we obtain the well-known probability of fixation of an allele with frequency *p* under drift and no selection.

We will now consider the *j*^*th*^ moments of the time until first passage, conditional on reaching *f*, which we denote as 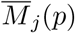. Importantly, note that 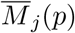 depends on *f*, but we have excluded *f* as a variable for simplicity of notation. According to these definitions, 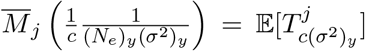 and 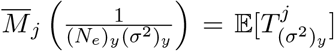 where it is implicit that 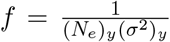 and 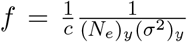 respectively. This relates the 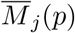 notation to the previous notation (Eqs. 29, 30, 33, 34). We will now compute the moments by first defining the useful quantity as done in [71]

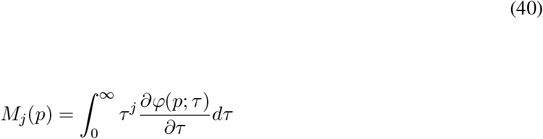

*M*_*j*_(*p*) is the unnormalized version of 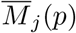 and differs by a factor *φ*(*p*). That is,

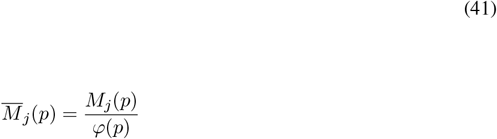

To find 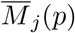, we will rewrite the Kolmogorov backwards equation and its boundary conditions in terms of *M*_*j*_(*p*), and then after solving the equation normalize *M*_*j*_(*p*) to obtain 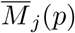. We begin with the first moment *M*_1_(*p*), and convert the backwards equation by differentiating both sides with respect to *τ*, multiplying by *τ*, and then integrating from 0 to ∞ to obtain:

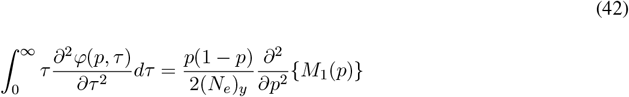

Through integration by parts, assuming 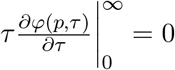 and noting *p < f* we obtain:

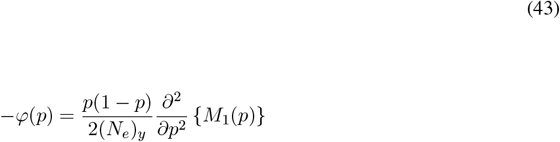

Substituting the ultimate probability of reaching *f*, 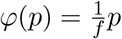, we obtain:

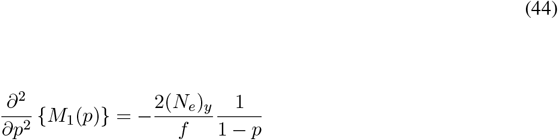

To derive the boundary conditions, we assume 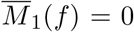 since the expected time to reach frequency *f* is 0 assuming *p* = *f*, and that 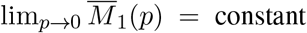. The second boundary condition reflects the scaling limit in Section 4 where the frequency distribution becomes invariant as the initial frequency vanishes. Multiplying by 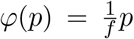 shows these boundary conditions are equivalent to the conditions *M*_1_(*f*) = 0, and *M*_1_(*p*) → *Kp* asymptotically as *p* → 0, where *K* is some constant (see Eq. 41). We can solve the ODE (Eq. 44) through integration:

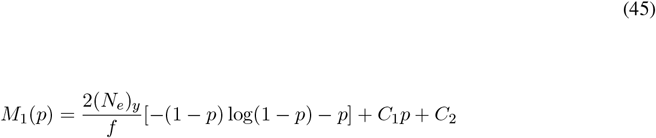

We first apply *M*_1_(*p*)− *> Kp* as *p* → 0, which is satisfied only if *C*_2_ = 0. This follows from the fact that the first term goes to 0 as 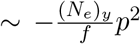 which can be seen from the Taylor expansion 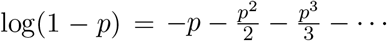. Applying the second boundary condition *M*_1_(*f*) = 0 we obtain:

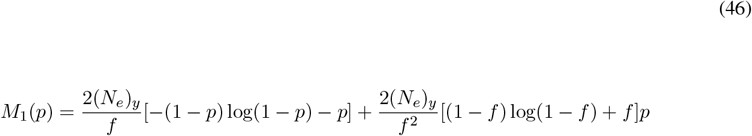

As a validation, normalizing (Eq. 46) by 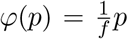 (Eq. 41) and then taking the limit as *f* → 1 we obtain:

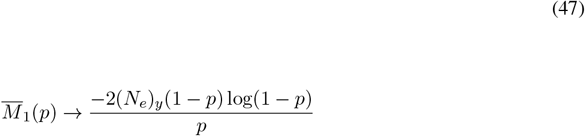

which is the mean time to fixation, found in [71] and previous sources but with a different approach. Next we will use the following Taylor expansion in (Eq. 46)

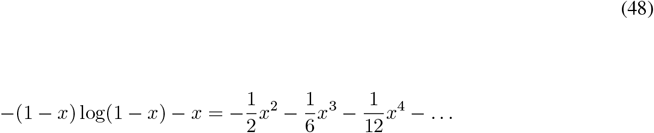

and assume *p <<* 1 and *f <<* 1 to let higher order terms vanish

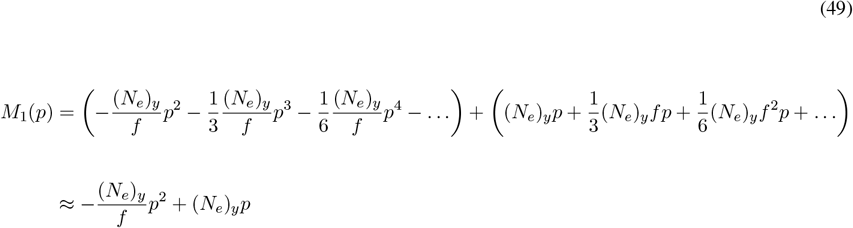

Normalizing by 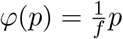 we obtain

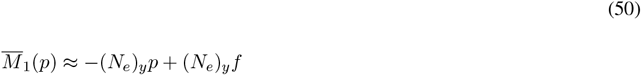

We will first consider the *c* ≥ 1 case and set 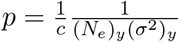 and 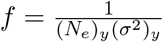 to obtain

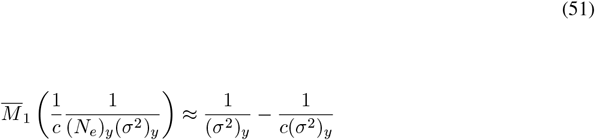

We can now use (Eq. 51) along with (Eq. 29) and the fact that 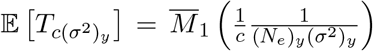 to obtain the expected offset in the posterior distribution:

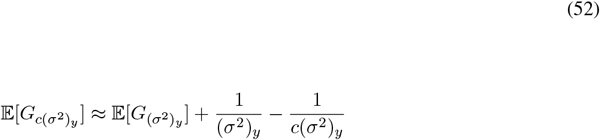

We will now take a step back to think about the result we have obtained. There are multiple but equivalent ways to interpret the equation, but we will suggest the following. First consider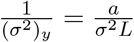 which is equal to the duration of a generation divided by the reproductive variance. Recall that the average population size during the drift phase is 1 + (*σ*^2^)_*y*_*τ* conditional on fixation, and therefore 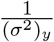 is the time it takes the population size to increase on average by one. We refer to this quantity as the effective duration of a generation, as an analogy to an effective population size, and use the notation Δ. The effective duration of a generation of a process can be thought of as the duration of a generation assigned to a Wright-Fisher model (holding *s*_*y*_, *g*_*y*_, (*N*_*e*_)_*y*_ fixed) that has overlapping trajectories of population size and the same distribution of coalescence times. If we let 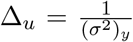 refer to the Δ of the underlying process, and 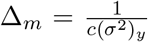 to the Δ of the model, we can obtain a cleaner expression

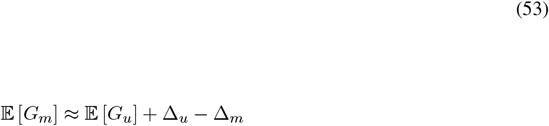

which holds for *c* ≥ 1. This change of notation in the sample distributions reflects the fact that the formula turns out to not depend on *c* explicitly. This is because we can also show that (Eq. 53) holds for 0 *< c <* 1 as well. To see this, we can revisit the offset formula for 0 *< c <* 1

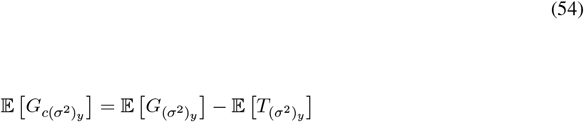

We can then go back to the expression for 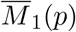 in (Eq. 50) and substitute the modified initial conditions 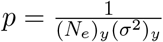 and 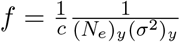 to obtain

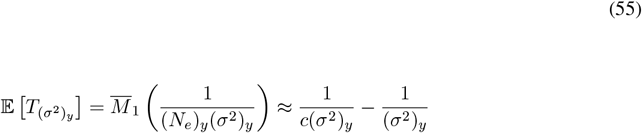

Substituting (Eq. 55) into (Eq. 54) we obtain

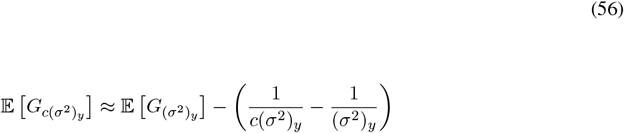

or equivalently

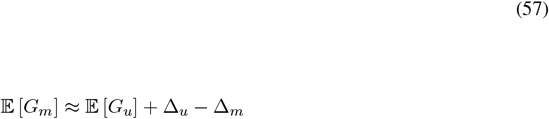

for 0 *< c <* 1, which is the exact same expression we obtained for *c* ≥ 1. It therefore follows that these formulas hold for all *c >* 0.

To obtain the offset in the variance under the parameter change, we will compute 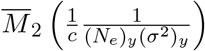. We can take a similar approach and differentiate the backwards equation with respect to *τ*, multiply by *τ* ^2^, and then integrate across all *τ* to obtain:

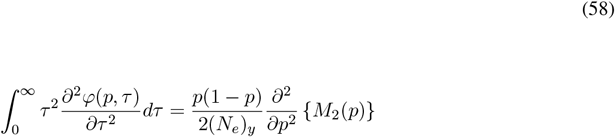

Using integration by parts, assuming 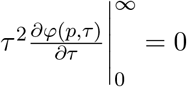, and rewriting the expression we obtain

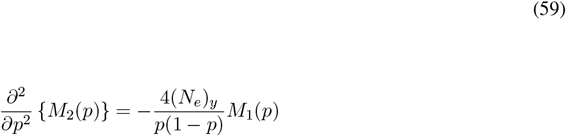

We must now substitute *M*_1_(*p*) into the equation. However, we are interested in an approximate solution under the assumption *p <<* 1 and *f <<* 1, so we will substitute (Eq. 49). To do the following derivation correctly, we must actually include higher order terms of *M*_1_(*p*) and show that they vanish in the final equation. To simplify the derivation shown here, we will not include those higher order terms and simply note that when we obtain the final answer those terms will show up as higher order products of *f* and *p* and vanish. We now substitute our approximation for *M*_1_(*p*):

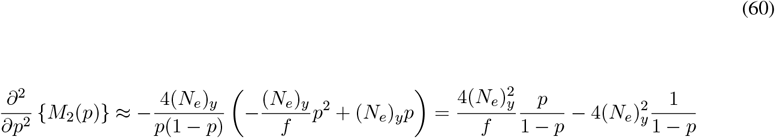

Integrating once we obtain:

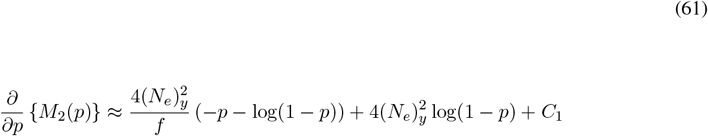

Using the following first order approximations, and again noting that higher order terms will vanish, we can substitute

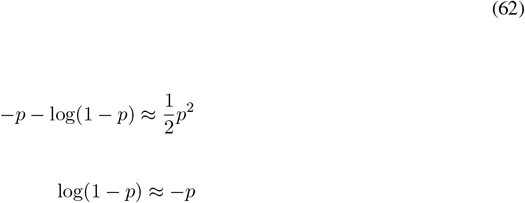

into (Eq. 61) to obtain

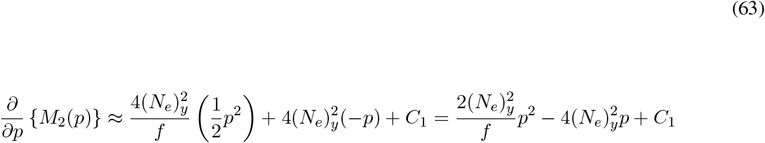

We then integrate a second time:

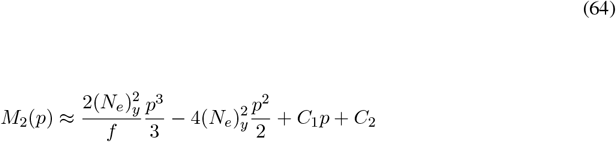

Similar to before, we can derive boundary conditions that reflect the scaling limit in Section 4, in particular, we assume 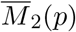 becomes constant, or equivalently *M*_2_(*p*) → *Kp* asymptotically, as *p* → 0. In addition, we assume 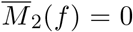 or equivalently *M*_2_(*f*) = 0. After applying both boundary conditions we obtain:

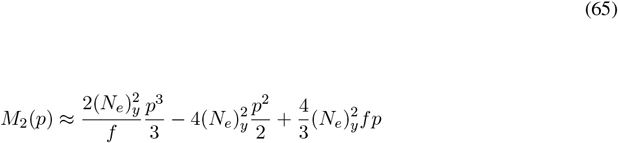

Now applying the normalization*φ*(*p*) we obtain:

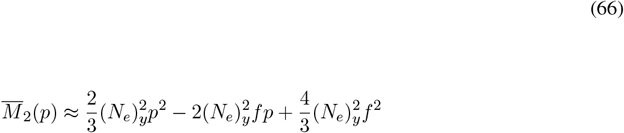

Lastly, substituting 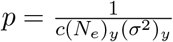 and 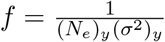 we obtain

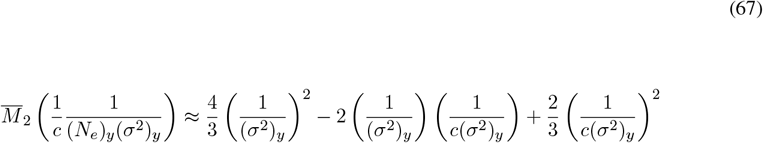

We can now compute the variance from the first two moments:

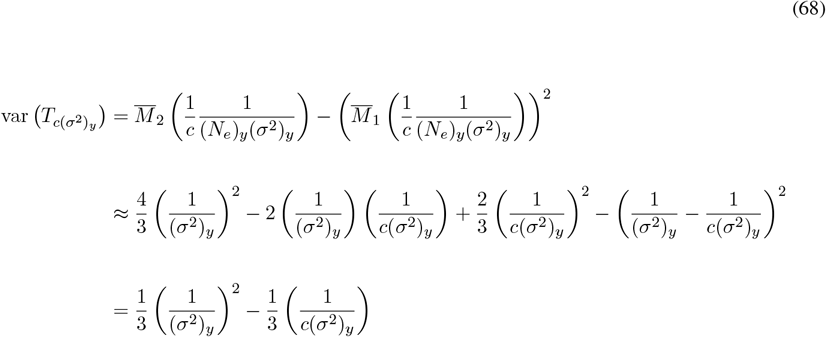

It therefore follows that the offset in the variance is:

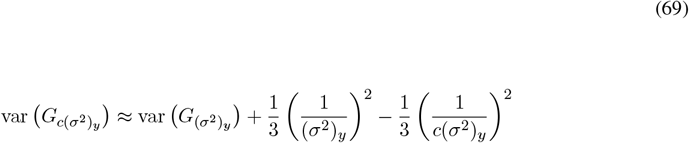

or equivalently

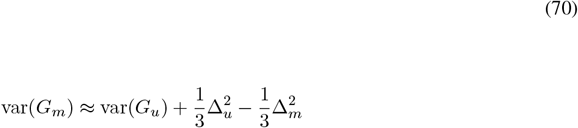

Although this derivation assumes *c* ≥ 1, we can do a similar calculation as for the mean using the variance offset formula for 0 *< c <* 1 to show the formula holds more generally for all *c >* 0.

## 8 Empirical Validation of Mean Offset Formulas Using Simulated Data

In the previous sections, we showed that the mean offsets in the posteriors of *g*_*y*_ and *s*_*y*_ when using a set of correct vs. incorrect parameter values are

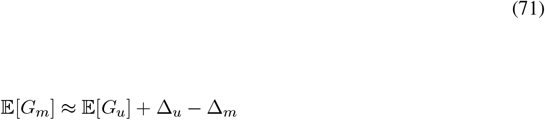

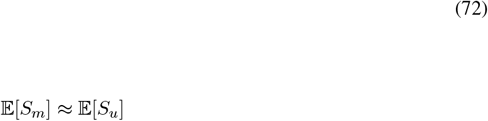

where Δ_*u*_, Δ_*m*_ are effective durations of a generation. *S*_*m*_ and *S*_*u*_ are random variables representing a sampled point from the posterior of *s*_*y*_. As stated before, the invariance of the inferred *s*_*y*_ (Eq. 72) is a direct consequence of the way trajectories are mapped between the scaled and unscaled processes, and therefore does not require an explicit calculation. Subscripts *m* and *u* denote whether the posterior is computed using the model parameters or the underlying process parameters, respectively.

To validate our theoretical results using simulations, we performed inference on simulated data and computed the mean offsets empirically. To do this, we generated a simulated data tree using the Wright-Fisher model with selection as defined above (more details can be found in [33]). The model parameters used were *s* = 0.06779, *g* = 200, *L* = 600, *N* = 10^9^, with the Wright-Fisher process conditioned on non-extinction at the final time point. A mutation rate of 5 per generation was used to assign the number of mutations, drawn from a Poisson distribution, to each branch of the tree. This sets the average tree length to 3000 mutations. 20 individuals at the final time-point were sampled, and the data tree was reconstructed as the coalescent tree of these individuals. The simulated tree used in our analysis is shown in (Fig. 1a).

**Figure 1:**
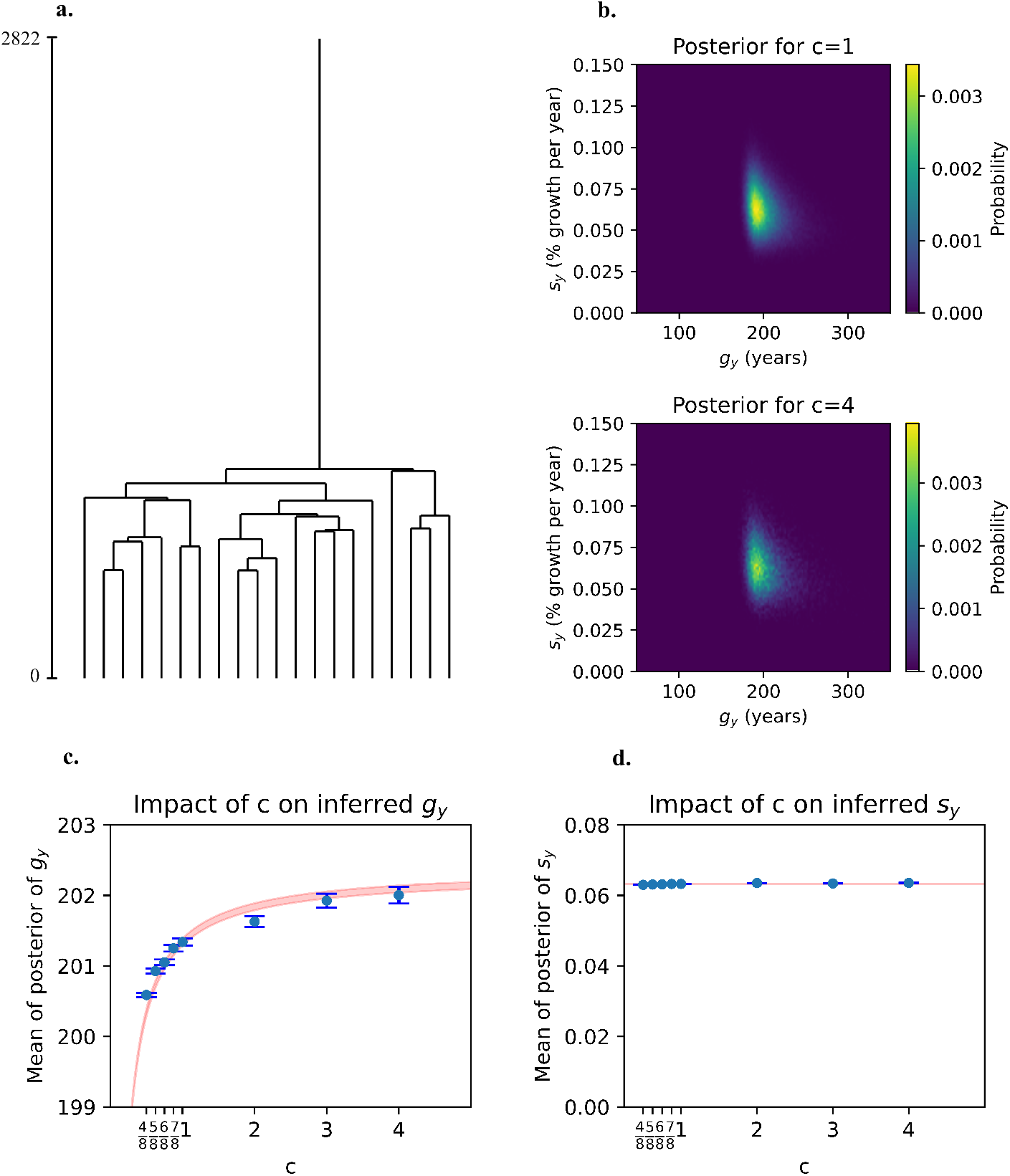
**a**. The simulated data tree, with the vertical axis representing the number of mutations accumulated along the branches. **b**. Two example posterior joint-distributions of *s*_*y*_ and *g*_*y*_ inferred from the data tree, using scaling factors of *c* = 1 and *c* = 4. **c**. The effect of varying the scaling factor *c* (which scales the model parameter *L*) on the inferred *g*_*y*_ posterior. Blue dots represent the numerically computed means of the posterior distributions, with error bars indicating the standard error due to finite sampling. The red shaded region shows the predicted theoretical offsets calculated using Equation 73, relative to the ground truth at *c* = 1 (where the model *L* matches the true number of generations used to generate the data tree). **d**. Identical to (c), but showing results for the computed means of the *s*_*y*_ posteriors and the corresponding theoretical offsets calculated using (Eq. 74).

Different posterior distributions were inferred from the data tree using the same Wright-Fisher model, a fixed *N* = 10^9^ that treats saturation as effectively infinite, and a fixed uniform prior on *s*_*y*_, *g*_*y*_ while varying the assumed number of generations in the model. The assumed number of generations for the different posteriors were *cL* for 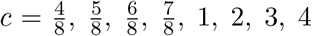,where *L* = 600. To superimpose the processes on an absolute time-scale, we assumed that each generation corresponds to one unit of time so that *a* = 600, or in other words 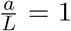.The mutation rate was set to 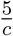 per generation to match the mutation rate per unit time in the data tree for all inferences.

Approximate Bayesian Computation (ABC) was used to obtain the posterior distributions. ABC simulates trees based on the specified priors and model, retaining parameter values that generate simulated trees similar to the observed data tree, as measured by a distance metric. These retained parameter values represent samples from the posterior distribution. Further details and software regarding ABC inference from coalescent trees, as used in our simulations, can be found in [33].

The number of data points sampled from each posterior corresponding to the different 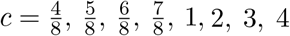 values were *n* = 322135, 247253, 201252, 172164, 149576, 64180, 36946, 25857 respectively. The empirical mean of the marginals for *g*_*y*_ and *s*_*y*_ were computed for each posterior, and the standard error of the means were computed through bootstrapping of the points sampled from the posterior with replacement.

We then derived the theoretical offsets of the mean. To do this, we computed the effective duration of a generation in the underlying process. Since 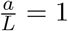 and 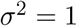 for the data tree, then Δ_*u*_ = 1. However, the model used for the inferences assumes *cL* number of generations with *σ*^2^ = 1, so 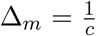.Therefore, the mean offset formulas are

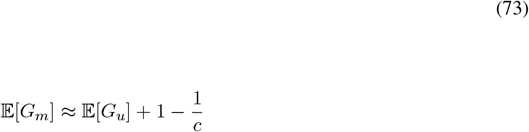

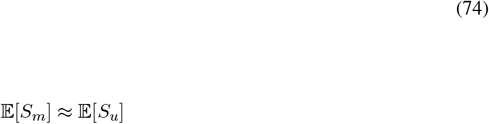

where 𝔼[*G*_*u*_] and 𝔼[*S*_*u*_] are the means of *G*_*u*_ and *S*_*u*_, respectively, estimated empirically using ABC with the correct parameter values. The equations predict the offsets from these estimated means.

We begin by plotting two example posterior distributions from the data tree using *c* = 1 and *c* = 4 (Fig. 1b, top and bottom, respectively). To precisely illustrate the results, we directly compare the empirical mean offsets with their expected theoretical values for all *c* (Fig. 1c). In this figure, the empirical mean calculated for each *c* is shown as a blue dot, with bootstrapped standard error of the mean represented by error bars. The red shaded region represents the theoretical offset calculated using (Eq. 73). More specifically, the upper and lower boundaries of this region are determined by setting 𝔼[*G*_*u*_] in (Eq. 73) to the upper and lower bounds of the error bar at *c* = 1, respectively. This provides a probable range for the true mean offsets. By definition, the red shaded region includes the blue dot at *c* = 1, as the offset should be zero when using the correct parameter values. (Fig. 1d) presents analogous results, but with blue dots representing the computed means for the posteriors of *s*_*y*_ and the red shaded region representing the theoretical offsets calculated using (Eq. 74). Our theoretical predictions of the mean offsets correctly match the simulated inferred values. We did not obtain simulated higher moments due to the computational challenge of resolving them numerically, a consequence of the exceedingly small predicted effect size.

## 9 General robustness for nonuniform prior distributions

Up to this point, we have derived explicit formulas for the offset of the mean and variance of the inferred posterior when varying (*σ*^2^)_*y*_ under the assumption of a uniform prior distribution. These formulas were derived from a set of general equations (final form of Eq. 26 and Eq. 27) which allow for arbitrary prior distributions. We will rewrite the general equations below. In particular, the offset of the mean is:

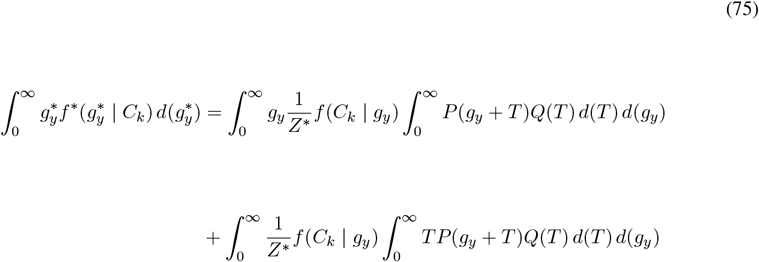

where

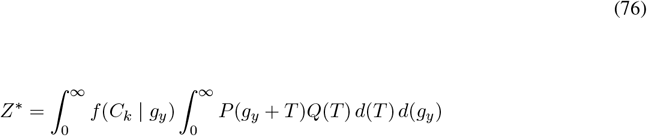

The offset of the second moment is given by:

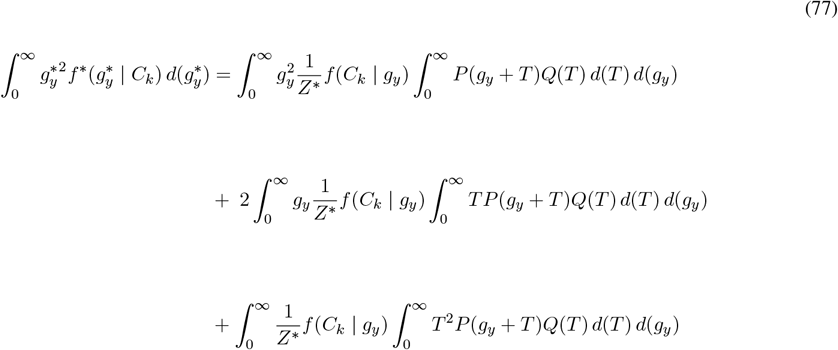

where *Z*^*^ is also computed from (Eq. 76). The offset in the variance is computed from both (Eq. 75) and (Eq. 77). Offset formulas for arbitrary *P* (*g*_*y*_) are then derived by substituting the explicit form of the prior into these equations. Although our goal here is not to create a catalog of equations under different priors, we will demonstrate the method’s utility with an example. We will consider one case of interest where we can explicitly compute the solution. In particular, we will assume an exponentially distributed prior 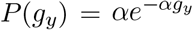,where *α >* 0. To begin, we will compute *Z*^*^ in terms of *Z* by substituting 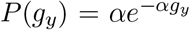 into (Eq. 76):

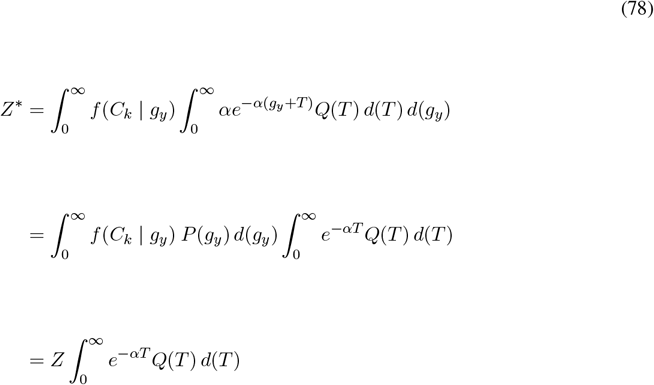

Substituting 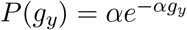 into (Eq. 75) we obtain:

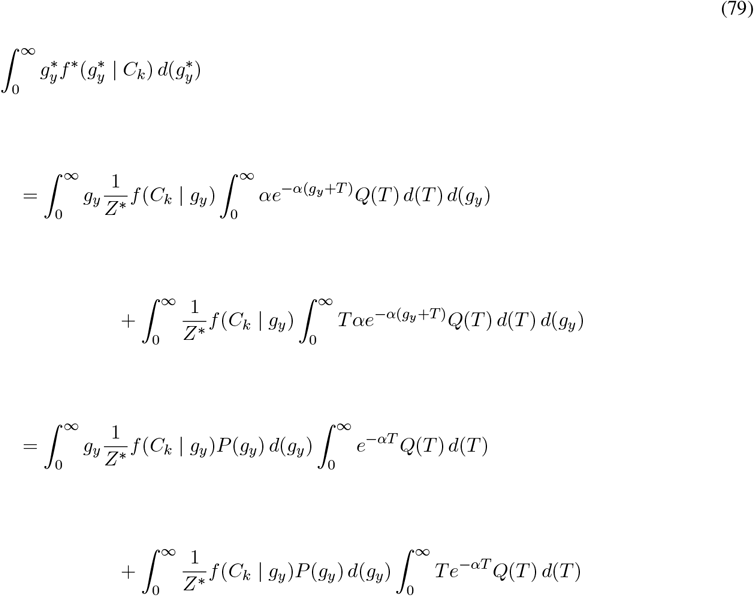

and then substituting (Eq. 78) into (Eq. 79) we obtain:

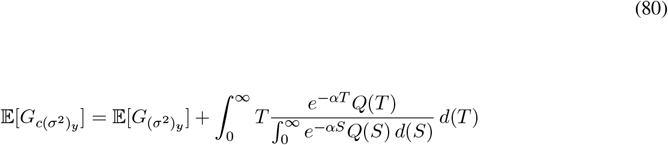

which holds for *c* ≥ 1. The argument of the integral in (Eq. 80), 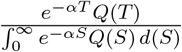 is a well-defined distribution, since it is non-negative and integrates to one. Therefore, 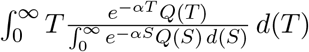 is an expectation. If we let 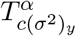 be a random variable denoting a sample from this distribution, then we can write:

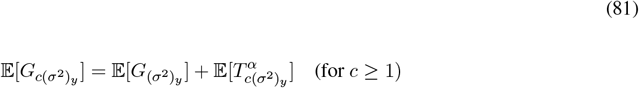

We can do a similar calculation for the 0 *< c <* 1 case to obtain:

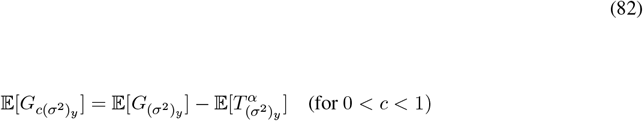

Similar to previous calculations, the subscripts *c*(*σ*^2^)_*y*_ or (*σ*^2^)_*y*_ for *T* ^*α*^ implicitly define both the starting frequency *p* and absorbing boundary *f* of the first passage time process of the corresponding *Q*(*T*). In particular, the *Q*(*T*) used to derive the expression for *c* ≥ 1 and for 0 *< c <* 1 are different. For *c* ≥ 1, 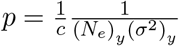 and 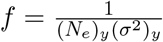,and 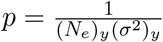 and 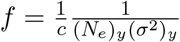 for 0 *< c <* 1.

A similar expression exists for the variance,

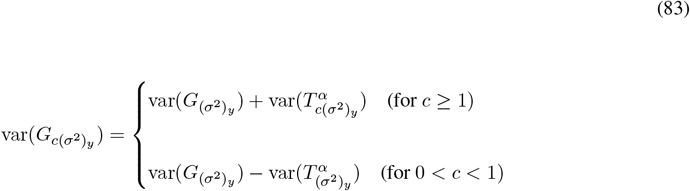

We can then compute the unknown first passage time dependent terms in (Eq. 81-82) and (Eq. 83) using the integral definition and then doing a Taylor expansion of the exponential factor:

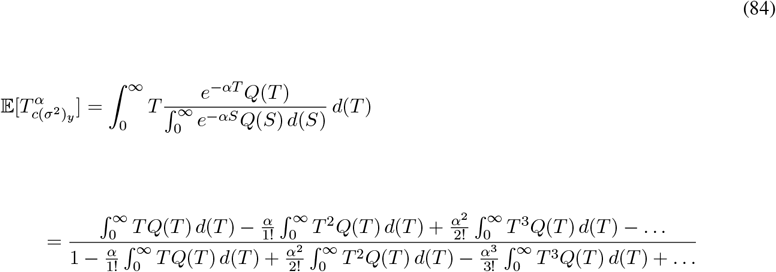

The exchange of the integral with the sum in (Eq. 84) is a valid operation and converges whenever 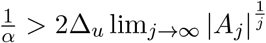 (see Supplement for details including Eq. 104 for definition of *A*_*j*_). The derivation of this result in the Supplement assumes *Q*(*T*) decays exponentially as *T* → ∞ asymptotically. Numerical simulations show 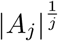 approaches a value in between 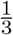 and 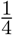 as *j* → ∞ (see Fig. 2 in the Supplement).

**Figure 2:**
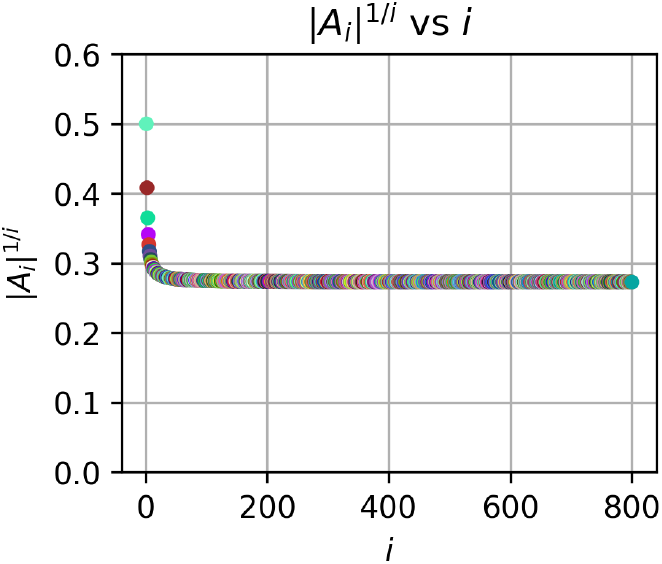
Different *A*_*i*_ for *i* = 1, 2, …, 800 are computed numerically and 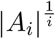 vs *i* is plotted. Different colors are used for each point for better visualization. The plot shows 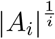 converging as *i* → ∞ which implies the magnitude of *A*_*i*_ decays exponentially as *i* increases.

The expectation in (Eq. 84) is now expressed as moments of *Q*(*T*) which are given by the Kolmogorov backwards equation. A similar Taylor expansion can be done for the case 0 *< c <* 1, except the exchange of the integral and sum is valid for 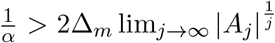, the difference being that Δ_*u*_ is replaced withΔ_*m*_.

In the Supplement, we derive the general form for the *j*^*th*^ moment of *Q*(*T*) in terms of effective generations using the Kolmogorov backwards equation. Using the 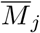 notation in (Eq. 84), we can write

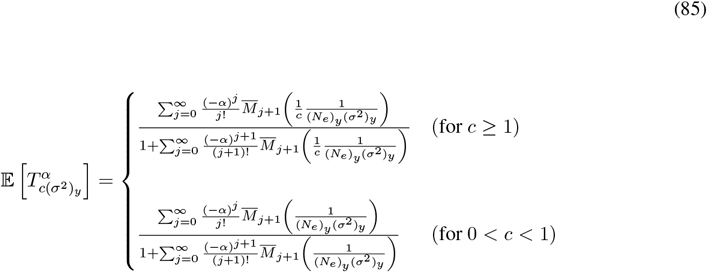

where the 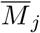 are given by (Eq. 104) in the Supplement. Substituting (Eq. 85) into (Eq. 81-82) we obtain:

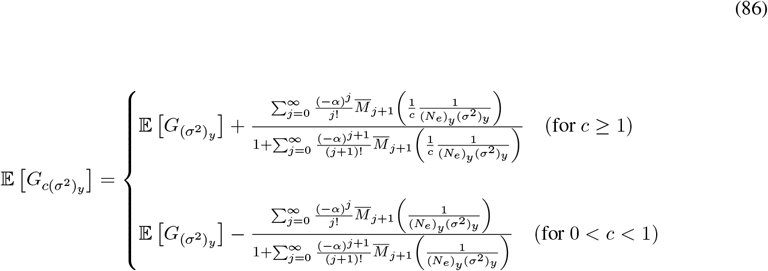

The unknown term in the variance offset can be computed in a similar manner through Taylor expansions of

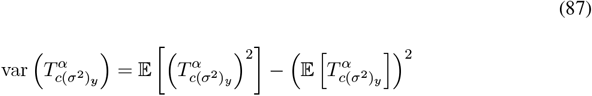

which then gives us:

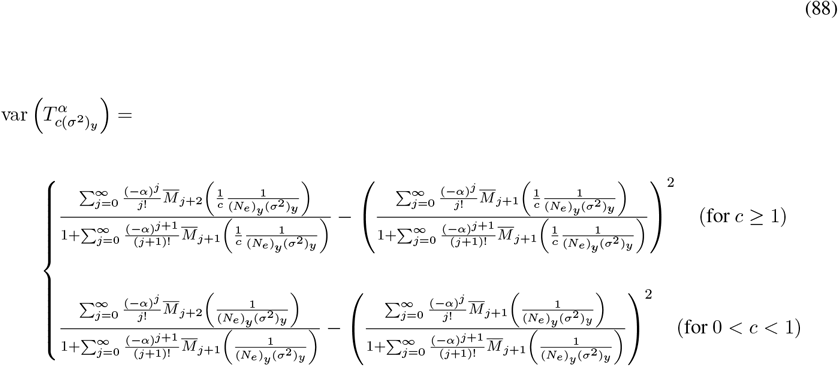

The offset of the variance is then obtained by substituting (Eq. 88) into (Eq. 83), and then substituting (Eq. 104) in the supplement for the final form. Notice that in the numerator of the first term of (Eq. 88), the moments have subscript *j* +2 as opposed to *j* +1. Lastly, similar to the uniform prior case, these expressions can be written as expansions of Δ. This is a general feature of these formulas since the conditions *p* and *f* can be written in terms of Δ.

Up to this point, we have considered two particular cases of prior distributions where the offset formulas can be solved explicitly: the uniform and exponential distribution. In both of these cases, the offset of the mean and variance of the posterior is impacted by an additive first passage time term. This occurs because the uniform and exponential distribution exhibit the properites *P* (*g*_*y*_ + *T*) = *P* (*g*_*y*_) and *P* (*g*_*y*_ + *T*) = *P* (*g*_*y*_)*P* (*T*) respectively. In the former case, we made this assumption explicitly while in the latter case implicitly. However, the uniform and exponential distribution are the only distributions that satisfy either of these properties, and so in general the relationship between the first passage time and the offsets of the posterior is not as simple. Although in general we cannot derive simple expressions, we can still compute the offsets numerically using the generalized equations.

One potential approach is to start by assuming *P* (*g*_*y*_) is a smooth and entire function, and use the Taylor expansion 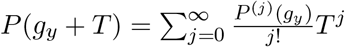 in (Eqs. 75-77) to obtain for case *c* ≥ 1:

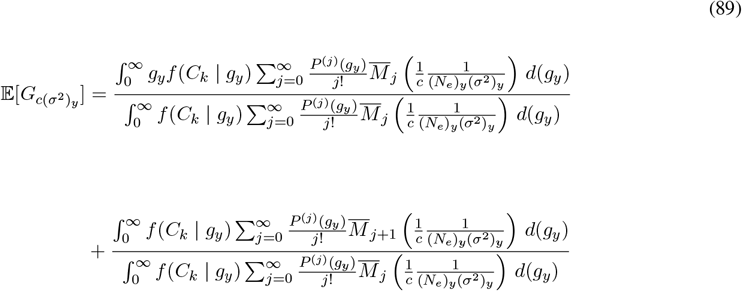

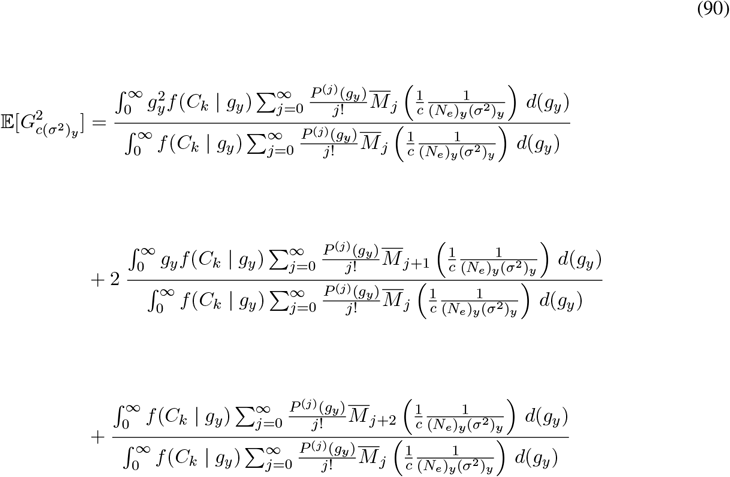

which show how the mean and second moment of the posterior change respectively. Special care should be taken to ensure that exchanging the integral and sum to obtain (Eqs. 89-90) is valid, such as establishing dominated convergence based on the properties of the assumed prior. The case 0 *< c <* 1 is more complicated because it not only requires using 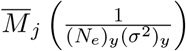 instead of 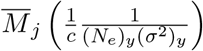 due to the different conditions on *p* and *f*, but also swapping 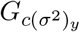 and 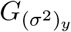 so that the left hand sides of (Eqs. 75-77) now refer to the expectation and second moment using the underlying (i.e. the unscaled) process. The posterior of the model process (i.e. the scaled process) is now convolved with the first passage time distribution requiring a transform or de-convolution process to invert the equation. We will not propose a method to solve for this case, and will instead leave it as an open problem.

(Eqs. 89-90) can be used to numerically approximate the offset formulas for arbitrary prior distributions under the appropriate assumptions given a sufficient number of sampled points from the inferred posterior.

This is possible because we can rewrite *f* (*C*_*k*_|*g*_*y*_) in terms of the posterior as 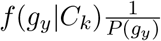,since the normalization constant for *C*_*k*_ shows up both in the numerator and denominator. Therefore, given *n* points sampled from the posterior, say 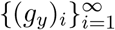,we can approximate the offset formulas by taking a weighted average over (*g*_*y*_)_*i*_ and 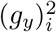 for the sample mean and second moment respectively using the following estimators:

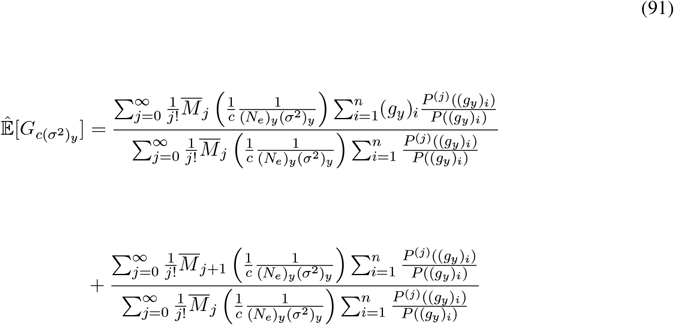

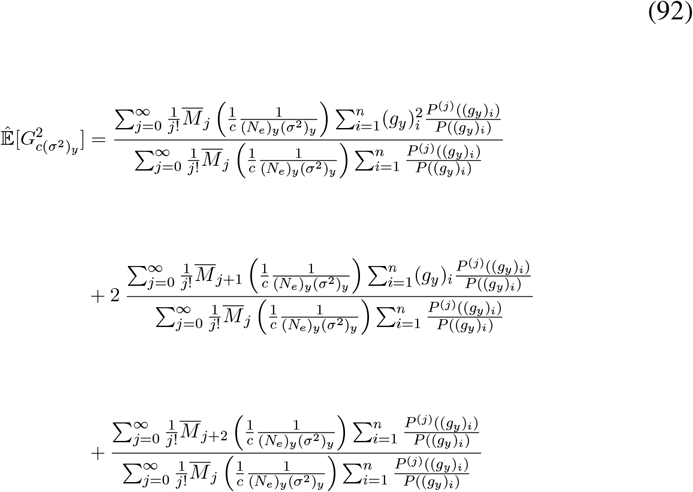

which hold for *c* ≥ 1. As before, the variance offset estimator is obtained by combining both equations (Eqs. 91-92). We can now use these estimators along with sampled points from the posterior distribution generated using any Bayesian inference framework (such as Approximate Bayesian Computation) with a non-uniform prior distribution and generate the offset of the mean and variance of the inferred *g*_*y*_ due to changing the model parameters.

As mentioned before, (Eqs. 91-92) can break down when the prior distribution is not entire, since integration to ∞ breaks down. In those cases, it may be possible to derive different estimators by centering the Taylor expansion around different points to obtain a larger radius of convergence, and then finite truncating the integral as an approximation. In addition, if dominated convergence cannot be established on the interval [0, ∞) and exchanging the integral with the sum is not justified, but these conditions can be established on compact intervals, then finite truncating the integral and letting the partial sums go to infinity could be a potential solution.

## 10 Discussion

We derived a set of general equations that describe how the posterior distribution of the inferred time of onset and fitness of a beneficial mutation change with the reproductive variance (*σ*^2^)_*y*_ (or equivalently the number of generations *L*) in Cannings models of population genetics. We also showed that the inferred posterior distribution of the selection coefficient in units of absolute time is independent of (*σ*^2^)_*y*_. Under the assumption of a uniform prior distribution, we derived the offset of the mean and variance of the posterior of the time of onset and related it to the moments of a first passage time distribution. We solved for the moments of first passage using the Kolmogorov backwards equation and related it to the effective duration of a generation 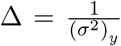.The derived formulas were validated with simulations. We then showed that the generalized equations can be used to derive formulas for arbitrary prior distributions as well, and as an example we considered the case of an exponentially distributed prior. In addition, we derived two estimators to compute the offset of the mean and variance numerically, but under restricted conditions and only for the case *c* ≥ 1 where the *c*(*σ*^2^)_*y*_ of the model process is larger than the (*σ*^2^)_*y*_ of the underlying process. All of these formulas were derived assuming 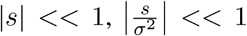 and *N >>* 1, and that 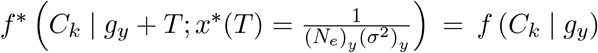 for all *T >* 0. If any of these assumptions break down significantly, then the derived formulas will not apply.

In terms of future directions, we make the following suggestions. In this work, we only focused on the offset of the mean and variance of the posterior. Extensions to higher moments are easily made by directly computing the offset of the *j*^*th*^ moment (i.e. the arbitrary *j*^*th*^ moment version of Eqs. 75-77) using a binomial expansion of (*g*_*y*_ + *T*)^*j*^ in the derivation to break apart the integral. Under a uniform prior assumption, the expression reduces further to products of the moments of the original posterior with a correct parameter set and moments of the time of first passage which are given by (Eq. 104) in the Supplement. It would also be interesting to check if there are simplified formulas for the centered moments as well, similar to the calculation for the variance. In addition, our framework allows for numerical simulations of the posterior offsets under arbitrary prior distributions using derived estimators (Eqs. 91-92) of the generalized equations (Eqs. 75-77). These estimators hold under restricted conditions and only for the *c* ≥ 1 case, but it may be possible to de-convolve the generalized equations or possibly (Eq. 22) for the 0 *< c <* 1 case. We suggest improving the numerical approach, clarifying the conditions under which the approximations hold, and developing a de-convolution method for the remaining case. In addition, the concepts introduced here could be extended to multiple competing or subclonal mutations, and it would be interesting to think about the limits where the equations or assumptions break down, for example, when multiple merger events occur. Lastly, our framework applies to other forms of data as well besides coalescent trees, such as time-course measurements of gene frequencies or site-frequency spectra, and so more rigorous calculations for other data forms should also be carried out.

In summary, we were able to develop a framework based on a first passage time approach to extend notions of robustness under an effective population size to inferences of selection and time of onset of mutations. This approach is not only useful conceptually, but can be used to correct inferences under changing model assumptions and new information using simple formulas instead of having to reanalyze data. As a result, this work furthers our understanding of the robustness of population genetics models with drift and selection in a way that can be directly applied to real-world data.

## 11 Acknowledgments

We acknowledge generous funding from the National Institutes of Health (NIH) –National Heart, Lung, and Blood Institute (NHLBI)– under grant R01HL158269 and its supplement, and the Leukemia & Lymphoma Society under grant 8041-24.

## 12 Supplement

The supplemental section will be divided into two parts. In the first part, we will establish a general formula for computing arbitrary *j*^*th*^ moments of the first passage time distribution. In the second part, we will justify the exchange of the integral with the sum in (Eq. 84) using Lebesgue’s dominated convergence theorem. In the derivation, we will make the assumption that *Q*(*T*) decays exponentially.

We will now derive the *j*^*th*^ moments of the first passage time distribution. The formula is obtained by solving the moment version of the Kolmgorov backwards equation inductively. To begin, we will consider the *j*^*th*^ moment transformed Kolmogorov backwards equation (similar to Eqs. 42 and 58 for the first two moments but we are instead multiplying by *τ* ^*j*^) and then assume *p <<* 1 so that the *p*^2^ terms vanish, or equivalently by replacing *p*(1 − *p*) with *p* to obtain the following approximation

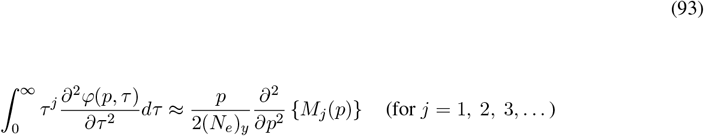

Recall that this equation is for computing the unnormalized moments. It turns out that solving the approximate form of the transformed equation (i.e. Eq. 93) is equivalent to solving the equation without the approximation and then letting the order *p*^2^ terms vanish in the solution. We will use this approximation because it simplifies the calculations. We will let the reader verify that both approaches are equivalent and produce the same answer.

Using integration by parts and assuming 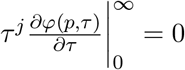 and *p < f*, we can rewrite (Eq. 93) as

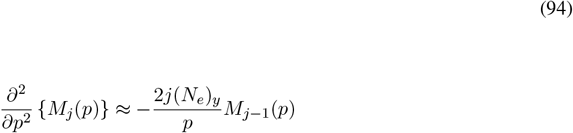

Here the boundary conditions are *M*_*j*_(*f*) = 0 and *M*_*j*_(*p*) = *Kp* asymptotically as *p* → 0. We claim that the general solution to the boundary value problem is

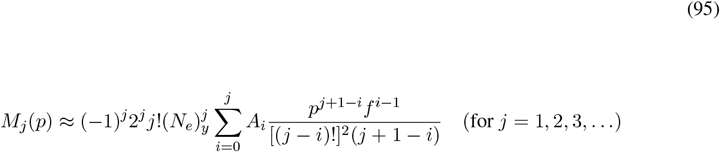

where 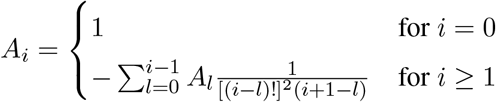

This answer was obtained by solving for the moments recursively for several iterations until we identified the pattern. For a rigorous proof, we will use induction.

We begin with the base case *j* = 1 and show

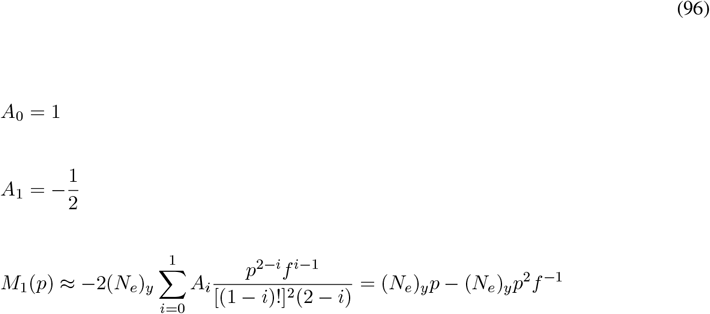

which matches (Eq. 49), and is hence the solution for *j* = 1.

For the inductive step we assume the formula is true for *M*_*j*_(*p*). Using this assumption in (Eq. 94) for *j* + 1 we can write

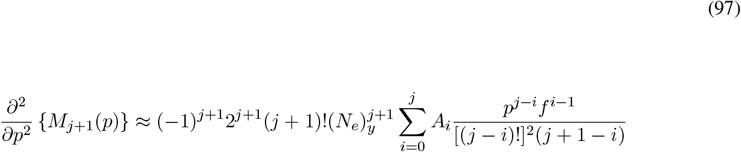

(Eq. 97) can then be integrated twice producing

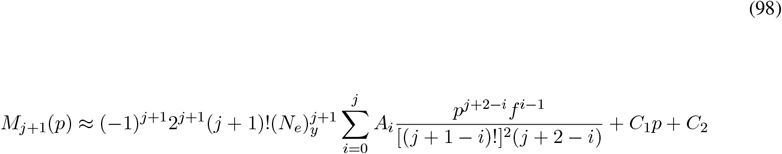

where *C*_1_ and *C*_2_ are constants of integration. *C*_2_ then vanishes after applying the asymptotic boundary condition

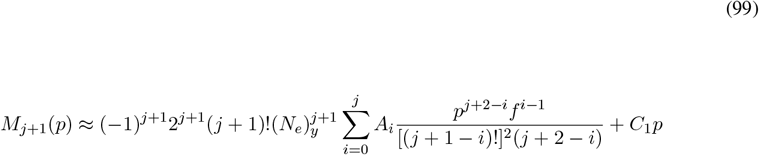

It will be useful to rewrite 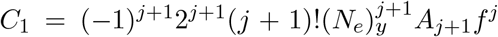, where *A*_*j*+1_ is an arbitrary constant up to this point

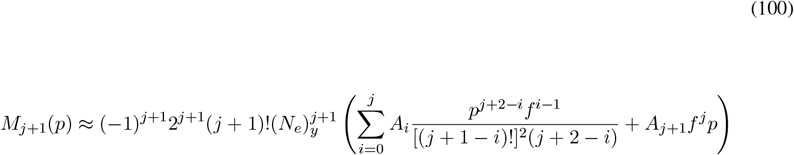

After applying the remaining boundary condition *M*_*j*+1_(*f*) = 0 we obtain

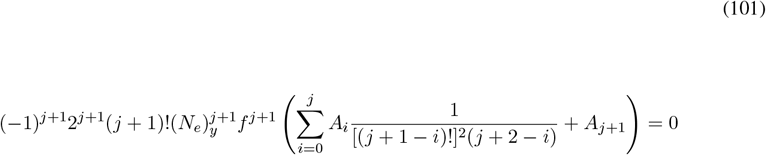

or in other words, 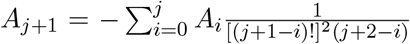. The result then follows because *A*_*j*+1_ has the same form as the *A*_*i*_ for *i* ≤ *j*, and since the *A*_*j*+1_ term can be absorbed into the sum and rewritten as

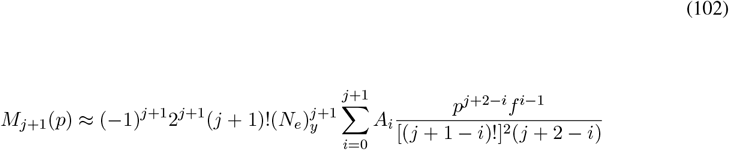

Now that we have established the formula for all *j* = 1, 2, 3, …, we can normalize the solution by 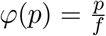 to obtain the moments of first passage

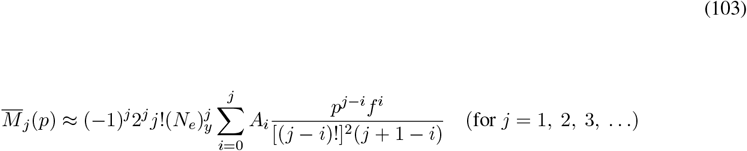

Lastly, we can substitute the appropriate *p* and *f* values. Recall that there are two cases (revisit section 7. Calculating the time of first passage). In particular, for *c* ≥ 1 we use 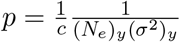 and 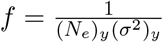, and for 0 *< c <* 1 we use 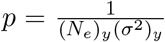 and 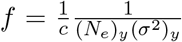.After substituting for both cases, and then using the definition of Δ_*m*_ and Δ_*u*_, we obtain the final form of the moments

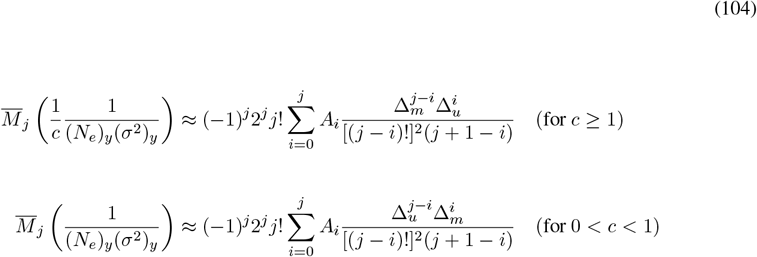

where 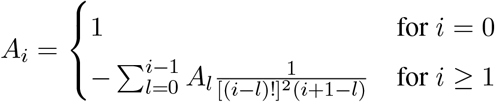

Simulated values of 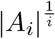 in (Fig. 2) show it is monotonically decreasing and converges to a value between 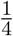 and 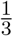. This implies *A*_*i*_ decays exponentially as *i* increases.

The remainder of the supplement will be used to show that the integral in (Eq. 84) can be exchanged with the sum. This can be done through Lebesgue’s dominated convergence theorem by showing that all of the partial sums before distributing the integral are bounded by some integrable function. However, we will first consider the asymptotic tail behavior of *Q*(*T*). In particular, we must argue that *Q*(*T*) decays sufficiently fast as *T* → ∞ so that the partial sums remain bounded.

We can see from (Eq. 104) that for the case *c* ≥ 1:

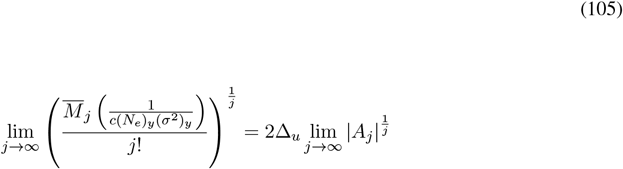

provided the limit exists (Fig. 2 shows the limit numerically and Fig. 3 shows Eq. 105). For the case 0 *< c <* 1, we simply replace Δ_*u*_ with Δ_*m*_ and 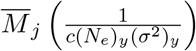 with 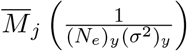 in (Eq.105). The existence of the limit (Eq. 105) suggests the density of first passage *Q*(*T*) asymptotically has an exponentially decaying tail with rate 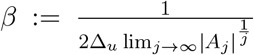, since the *j*^*th*^ moments of *Q*(*T*) match the moments of the exponential distribution in the limit. Although a rigorous proof of this result should be established using large deviation theory, to make progress we will take as an assumption that *Q*(*T*) is asymptotically exponential for the remainder of the supplement. Note that we will also use the notation *β* for the remainder of the supplement.

**Figure 3:**
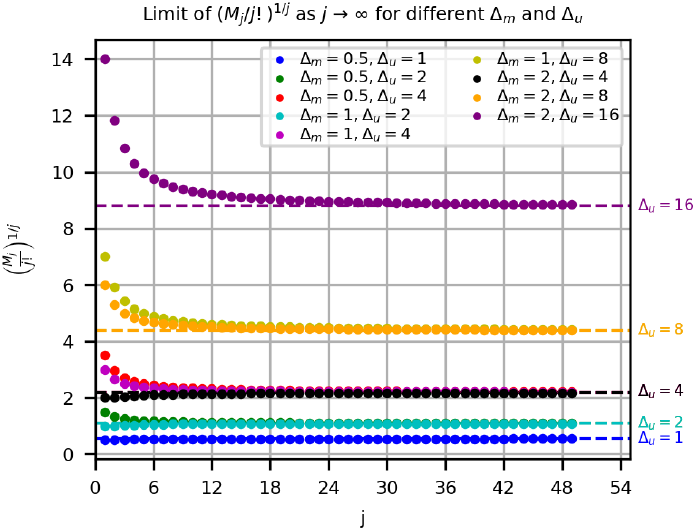
The limit in (Eq. 105) is shown numerically by computing 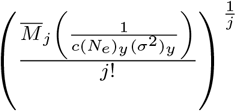 for *j* = 1, 2, 3, …, 50 using the definitions provided in (Eq. 104) for different parameter choices of Δ_*m*_ and Δ_*u*_. The simulated values are plotted against *j* for each parameter choice all superimposed. Δ_*u*_ is always kept larger than Δ_*m*_ so that the condition *c* ≥ 1 is satisfied. The choices of Δ_*m*_ and Δ_*u*_ are shown in the figure legend with a corresponding dot color that identifies each parameter choice with a plot. The theoretical limit (right hand side of Eq. 105), is approximated numerically as 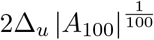 for each plot using the definition provided in (Eq. 104) and are superimposed as horizontal dotted lines. The theoretical limits depend only on Δ_*u*_, and are labeled to the right of their corresponding horizontal line according to the range of different Δ_*u*_ used. As expected, the plotted curves converge to their theoretical limits independent of the Δ_*m*_ used showing that (Eq. 105) holds.

We will now consider the partial sums of (Eq. 84). It is easy to show with the triangle inequality that

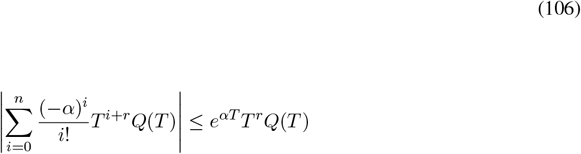

where *r* is a non-negative integer. Next we show the right hand side of (Eq. 106) is an integrable function. We consider the integral of (Eq. 106) and rewrite as follows

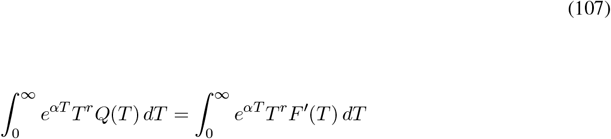

where *F* (*T*) is the CDF of *Q*(*T*). Using *F* ^*′*^(*T*) = (1 −*P* (*T* ^*RV*^ ≥ *T*))^*′*^ = −*P*^*′*^(*T* ^*RV*^ ≥ *T*), where *T* ^*RV*^ is the random variable with distribution *Q*(*T*), we obtain

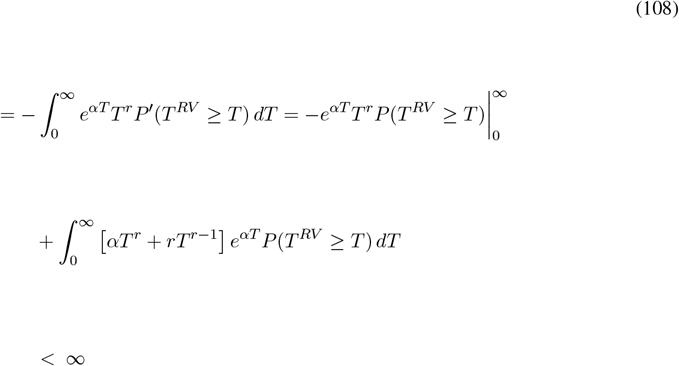

which follows from the assumption that *Q*(*T*) is asymptotically exponential, that is that 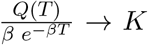 (*K* is constant) as *T* → ∞, so long as *α < β*. Under this assumption, dominated convergence is satisfied and the exchanging of the integral and sum in (Eq. 84) is justified.

